# Eosinophil-epithelial interactions mediate protective intestinal remodeling during food allergy

**DOI:** 10.64898/2026.06.04.730224

**Authors:** Patrick W. Darcy, Vitoria M. Olyntho, Albana Kodra, Zachary Kerner, Maria C.C. Canesso, Sandra Nakandakari-Higa, Gabriel Victora, Daniel Mucida

**Author notes:** Current address: Department of Cell Biology, Albert Einstein College of Medicine, New York, USA. Correspondence (P.W.D.); (D.M.).

## Abstract

Food allergies are associated with progressive gastrointestinal symptoms driven by exacerbated mucosal type-2 immunity. Here, we investigated whether cellular interactions between the gut epithelium and innate immune cells regulate the severity of allergic symptoms in mice. Using the BALB/c OVA-alum food allergy model, we observed that repeated oral allergen challenges remodel the gut epithelium, expanding tuft cells, while shifting the intestinal stem cell niche toward a fetal-like repair state. Using uLIPSTIC, we systematically characterized in vivo immune-epithelial interactions and found that eosinophils and mast cells directly interact with intestinal epithelial cells (iECs) in an allergen challenge-dependent manner. Epithelial subset-specific uLIPSTIC provided further resolution and revealed that eosinophils and mast cells contact enteroendocrine cells and Paneth cells in a regionally compartmentalized manner. Allergic challenge was associated with rapid eosinophil migration towards the crypts and modulation of iEC differentiation. Depletion of eosinophils using two independent approaches reversed key markers of food allergy-associated epithelial remodeling, while exacerbating allergic diarrhea and mortality from anaphylactic shock. These findings establish eosinophils as orchestrators of protective epithelial remodeling in food allergies.

## Introduction

In the gut, type 2 immunity acts as a biological quality control system that simultaneously promotes nutrient acquisition from food while limiting damage caused by helminths, venoms, and xenobiotics (Florsheim et al., 2021; Palm et al., 2012). These diverse noxious stimuli trigger conserved effector mechanisms, such as proteases, lipid mediators, mucus secretion, and modulation of gut motility, which act in concert to neutralize and expel the offending agent (Palm et al., 2012). Within this conceptual framework, food allergies arise when type 2 immune effector pathways are inappropriately activated by innocuous dietary antigens, reflecting a miscalibration of the quality control system. The cause of this breakdown is likely multimodal, with genetic makeup, early life antigen exposure, maternal allergy status and diet, and environmental factors contributing to food allergy sensitization (Du Toit et al., 2024; Florsheim et al., 2021; Perkin et al., 2016; Stein et al., 2016). Despite the clinical burden of food allergies, physical crosstalk between epithelial and immune cells remains largely uncharacterized, and the mechanisms by which such interactions shape allergic responses in the gut are unknown.

The intestinal epithelium is central to the miscalibration of quality control in food allergies, serving as both the primary site of allergen exposure and the source of alarmins that initiate type 2 immunity. Epithelial permeability varies across inbred mouse strains and correlates with food allergy susceptibility (Gertie et al., 2022; Hoyt et al., 2025). In addition, repeated oral allergen challenges disrupt tight junctions, allowing greater quantities of food antigens to cross the barrier (Yamani et al., 2021). Beyond its barrier function, the epithelium actively senses noxious stimuli and engages conserved type 2 effector responses, including “weep-and-sweep,” a coupling of fluid secretion into the gut lumen and smooth muscle contraction (Kopp et al., 2023; Perniss and Bankova, 2024). Tuft cells are a central node in this effector mechanism, producing acetylcholine that coordinates epithelial cell fluid secretion (Billipp et al., 2024; Ndjim et al., 2024; Touhara et al., 2026). Tuft cells also secrete the alarmin IL-25, which activates ILC2s and engages a feedforward loop of epithelial remodeling, characterized by tuft cell expansion (Gerbe et al., 2016; Howitt et al., 2016; von Moltke et al., 2016). Allergic diarrhea shares key features of weep and sweep, including fluid secretion and accelerated transit. However, the extent to which food allergies elicit epithelial remodeling beyond tuft cells remains unknown. Such remodeling would require the re-specification of intestinal stem cell (ISC) progeny, implicating the ISC niche as a target of allergen-driven immune-epithelial crosstalk.

Eosinophils are constitutively present in the intestinal lamina propria, whereas mast cells are relatively rare in the steady-state gut (Galli et al., 2020; Mishra et al., 1999). During type 2 inflammation, ILC2s coordinate additional eosinophil recruitment to the gut, whereas mast cells undergo pronounced expansion and accumulate near the epithelium (Arifuzzaman et al., 2022; Bachtel et al., 2025; Jacobsen et al., 2021; Xenakis et al., 2018). These two populations are activated through distinct mechanisms: mast cells are engaged primarily by IgE-Fcχr1 crosslinking, whereas eosinophils are stimulated by soluble factors, including epithelial alarmins and cytokines such as IL-5 and CCL11 (Bachtel et al., 2025; Jacobsen et al., 2021). Mast cells are required for anaphylaxis, expulsive diarrhea, and conditioned avoidance behavior, establishing their central role in orchestrating intestinal allergic responses (Bachtel et al., 2025; Brandt et al., 2003; Florsheim et al., 2023; Plum et al., 2023). In contrast, the function of eosinophils in food allergies is poorly understood.

Given the relevance of immune and intestinal epithelial cells (iECs) in food allergy, understanding the interactions between these compartments is critical for deciphering the development and chronicity of food allergy. Although iEC-mast cell interactions have been described, existing knowledge is largely restricted to communication at a distance via the secretion of soluble factors (Bachtel et al., 2025; Galli et al., 2020). Contact-dependent interactions between epithelial and immune cells remain largely unexplored, given the absence of tools capable of molecularly recording physical cell-cell contacts *in vivo* with cellular resolution. Universal (u)LIPSTIC enables sortase-mediated covalent labeling of immune cells that physically interact with epithelial cells expressing FLAG-tagged sortase enzymes (Nakandakari-Higa et al., 2024; Pasqual et al., 2018). Using Villin (all iEC)– and iEC subset-specific uLIPSTIC approaches, we comprehensively mapped iEC-immune interactions in the steady state and food allergy. We found that repeated oral food allergen challenges elicited progressive mast cell-iEC and eosinophil-iEC interactions, which were paralleled by epithelial remodeling, including tuft cell expansion and reprogramming of the intestinal stem cell niche. Organoid-eosinophil co-culture coupled with *in vivo* eosinophil depletion approaches revealed that eosinophil-iEC interactions regulate allergic epithelial remodeling. In addition, eosinophil depletion exacerbated allergic diarrhea and increased mortality from anaphylactic shock. Collectively, these experiments established eosinophils as critical cellular effectors that protect the host through direct engagement with the intestinal epithelium, functioning as a cellular arm of the gut quality control system.

## Results

### The Intestinal epithelium is remodeled in food allergy

Protective type 2 immune responses, elicited by either helminth infection or protist colonization, induce remodeling of the intestinal epithelium, including a rapid increase in the number of tuft and goblet cells (Gerbe et al., 2016; Howitt et al., 2016; von Moltke et al., 2016). To characterize the changes in epithelial composition during food allergy, we employed the BALB/c OVA-alum food allergy model (**Figure 1A-top**) (Brandt et al., 2003). With repeated OVA oral challenges (OVA-OC), OVA-sensitized mice developed progressively worsening allergic diarrhea (**Figure 1A-bottom**).

**Figure 1.**
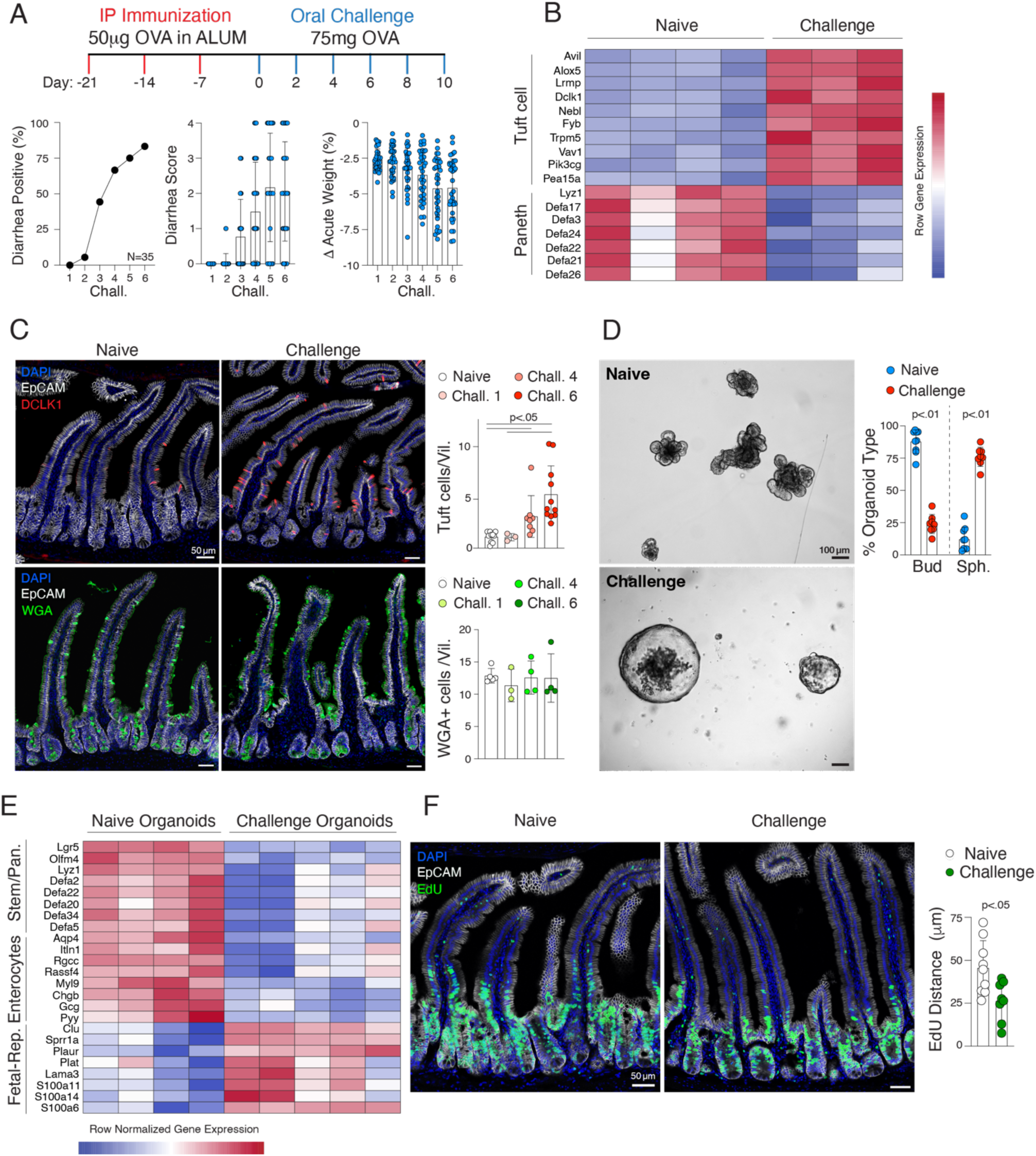
Epithelial remodeling in food allergy. **A**. Scheme for the food allergy model used throughout the study. BALB/c mice were intraperitoneally immunized weekly with OVA-Alum, followed by six OVA gavages at ten-day intervals. After each oral OVA challenge, the mice were observed for 2 h, and the incidence (bottom left) and severity (bottom middle) of diarrhea were scored. The variation in body weight before and 2 h after the challenge was assessed (bottom right; n=35). **B**. Bulk RNA-sequencing analysis of FACS-sorted EpCAM^+^ small intestine epithelial cells isolated from oral challenge naïve (n=4) and oral challenge experienced (48 h after third OVA-OC) (n=3) mice. All genes displayed are significantly differentially expressed between conditions, Wald test with Benjamini–Hochberg FDR correction. **C**. Representative images of tuft cells (DAPI-blue, EpCAM-White, Dclk1-Red) (top) and goblet cells (DAPI-blue, EpCAM-white, WGA-green) (bottom) in the duodenum of challenge-naïve (left) and challenge-experienced (3 h post-challenge 6) (middle) mice. Quantification of tuft cells (right, top) and goblet cells (right, bottom) per villus at the indicated time points (n=3-12). Ten villi per mouse were counted. Data were log10 transformed, and one-way ANOVA with Tukey’s multiple comparison correction was performed. **D-E**. Organoids derived from the small intestines of challenge-naïve mice (n=9) (top) and challenge-experienced mice (at least four OVA-OC) (bottom) (n=8). **D**. Organoids were quantified based on their morphology using Welch’s T test. **E.** Bulk RNA-sequencing was performed (challenge naïve, n=4; challenge experienced, n=5). All genes displayed are significantly differentially expressed between conditions, Wald test with Benjamini–Hochberg FDR correction. **F.** Representative images of duodenum from challenge naïve mice (n=9) and challenge experienced mice (3 h post sixth OVA-OC) (n=8), 17 h post EdU administration. The distance traveled by EdU above the crypt-villus transition point was quantified in micrometers (μm), and Welch’s T test was performed. DAPI-Blue, EpCAM-White, EdU-Green. Each dot in the bar graphs represents a biological replicate derived from a unique mouse, and the error bars represent one standard deviation from the mean. In the heatmaps, each column represents a biological replicate derived from a unique mouse.

To track epithelial remodeling in food allergy, bulk RNA sequencing was performed on FACS-purified small intestine epithelial cells from sensitized mice either naïve to OVA-OC (OVA-OC-naïve) or after three OVA-OCs (OVA-OC-experienced). Using the Broad Institute’s single-cell atlas of gut epithelial cells as a reference for epithelial cell gene signatures (Haber et al., 2017), we observed that most tuft cell-specific genes (69 of 97), but only a small fraction of goblet cell-specific genes (10 of 84), were enriched in OVA-OC-experienced mice (**Figure 1B, Data Table 1**). Immunofluorescence imaging confirmed that the number of tuft cells per villus progressively increased with repeated OVA-OC exposure, whereas the number of goblet cells remained relatively stable (**Figure 1C**), demonstrating that food allergy selectively expands tuft cells but not goblet cells, a pattern distinct from helminth infection. Of the nine Paneth cell-specific genes detected, seven were significantly suppressed in OVA-OC-experienced mice, suggesting reduced Paneth cell function upon repeated challenges (**Figure 1B, Data Table 1**).

Given that Paneth cells regulate the intestinal stem cell niche, we next addressed whether this suppression had functional consequences by generating intestinal organoids from OVA-OC-naïve and OVA-OC-experienced mice. Organoids derived from OVA-OC-experienced mice failed to bud and instead displayed a spheroid morphology. In contrast, crypts isolated from OVA-OC-naïve mice formed organoids with a conventional budded morphology (**Figure 1D**). Organoids derived from the fetal or injured gut have previously been observed to grow as spheroids, and this growth pattern is associated with the induction of a “fetal-repair” program in the stem cell niche (Fordham et al., 2013; Mustata et al., 2013; Sato et al., 2009). Bulk RNA sequencing confirmed that organoids from OVA-OC-experienced mice expressed the canonical spheroid-associated gene expression program, which included significantly higher expression of fetal-repair-associated genes (*Clu, Lama3*) and low expression of Paneth cell (*Lyz1, Defa2*), conventional stem cell (*Lgr5, Olfm4*), and mature enterocyte (*Aqp4, Itln1*) genes (**Figure 1E; Data Table 2**). *In vivo* EdU pulse-labeling revealed that epithelial cells showed reduced migration along the villus axis in OVA-OC-experienced mice, consistent with diminished proliferation and suppressed stem cell output (**Figure 1F**). Collectively, these data demonstrate that repeated oral allergen challenges remodel the cellular composition of the intestinal epithelium and reprogram the functional properties of the stem cell niche.

### Physic al interactions between iECs and immune cells in food allergy

Helminth– and protist-driven epithelial remodeling requires coordinated immune-epithelial crosstalk, including IL-4 and IL-13 signaling (Gerbe et al., 2016; Howitt et al., 2016; von Moltke et al., 2016). Physical interactions may also contribute, but the lack of appropriate tools has prevented direct assessment. We hypothesized that analogous immune-epithelial interactions mediate food allergy-associated epithelial remodeling. To identify such interactions, we used uLIPSTIC, a mouse genetic tool that records physical cell-cell interactions *in vivo* (**Figures 2A, S1A**) (Nakandakari-Higa et al., 2024). In Villin-uLIPSTIC mice, intestinal epithelial cells express a FLAG-tagged, cell membrane-anchored sortase enzyme (**Figure S1B**). When a biotin-substrate-loaded donor epithelial cell contacts an immune cell expressing the substrate-acceptor G5-Thy1.1, biotin is covalently transferred to the immune cell, labeling the interacting cells (**Figure 2A**). We compared immune cell labeling between sensitized mice that were naïve to OVA-OC and those with active diarrhea following the sixth OVA-OC.

**Figure 2.**
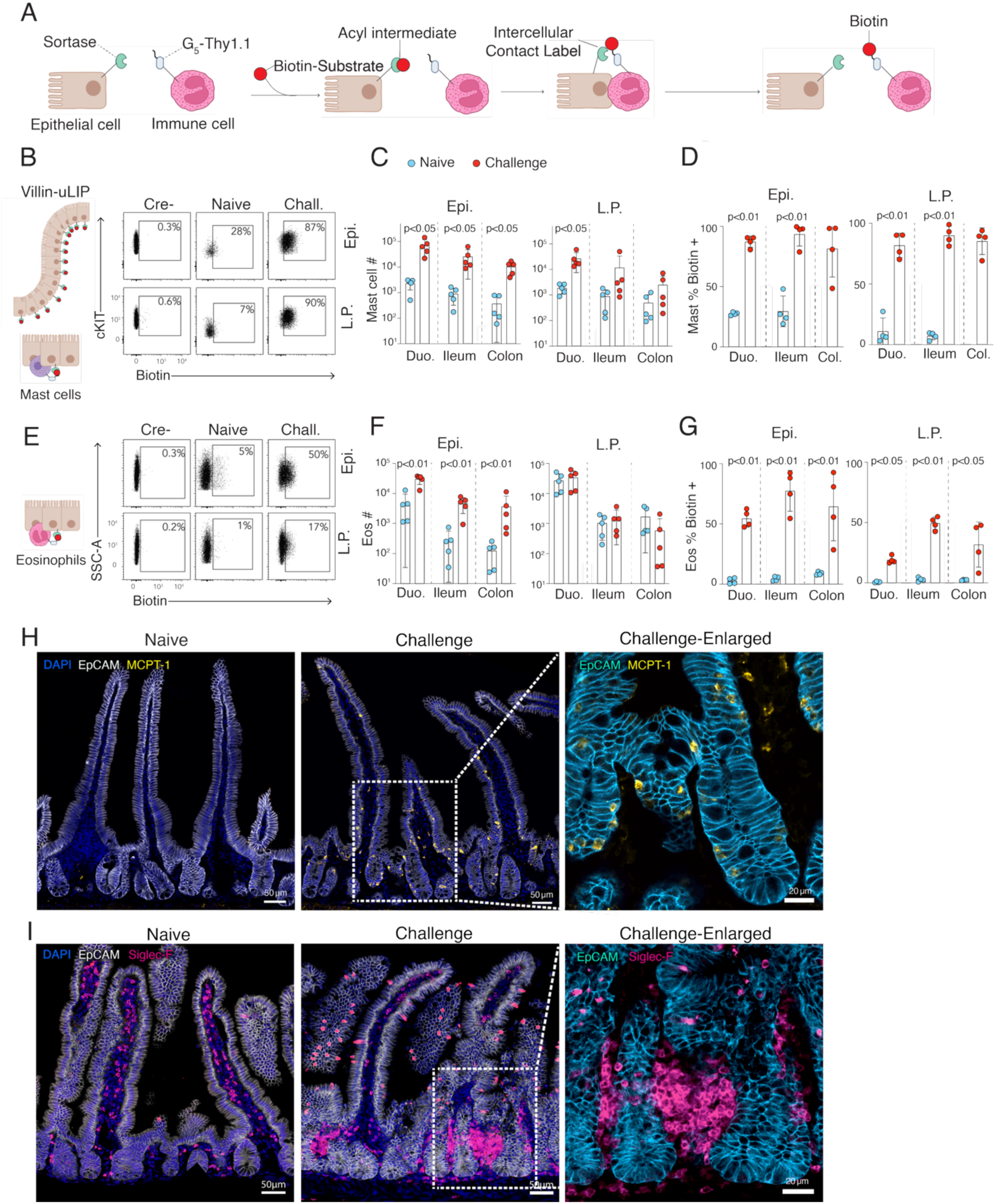
Eosinophils and mast cells engage in allergen-dependent interactions with epithelial cells. **A**. Scheme for the Villin-uLIPSTIC approach: Epithelial cells expressing sortase covalently transfer LPETG-biotin substrate to physically interacting immune cells, enabling the identification of epithelial-contacting immune cell populations. **B-G**. Flow cytometry analysis of Villin-uLIPSTIC biotin labeling in mast cells (**B-D**) (CD45+, TCRγδ-, TCRβ-, cKIT+, Fcer1+) and eosinophils (**E-G**) (CD45+, TCRγδ-, TCRβ-, CD11b+, SiglecF+, SSC-A^hi^). **B, E**. Representative flow cytometry plots from Cre– uLIPSTIC +/+ control mice after sixth OVA-OC, Villin-CreERT2+ uLIPSTIC +/+ mice challenge naïve, and Villin-CreERT2+ uLIPSTIC +/+ mice after sixth OVA-OC, stratified by epithelial location (epithelial-associated, top; lamina propria, bottom). **C, F**. Quantification of absolute immune cell numbers in challenge naïve and sixth OVA-OC mice. **D, G**. Quantification of the percentage of the indicated immune cell types labeled with biotin in challenge naïve and sixth OVA-OC mice. Welch’s T test or one-way ANOVA with Tukey’s multiple comparisons correction was used where applicable. **H-I**. Confocal imaging of the small intestine duodenum from challenge-naïve mice and 3 h after the sixth OVA-OC. **H**. Mast cells (DAPI-dark blue, EpCAM-white, Mcpt1-yellow). **I**. Eosinophils (DAPI-dark blue, EpCAM-white, SiglecF-magenta). Note that tuft cells are EpCAM+, SiglecF+, and eosinophils are EpCAM-, SiglecF+. Each dot in the bar graphs represents a biological replicate derived from a unique mouse, and the error bars represent one standard deviation from the mean.

We first quantified iEC-labelling of intraepithelial lymphocytes (IELs), a population of immune cells residing in the epithelial layer (Hoytema van Konijnenburg et al., 2017). Across IEL subsets, no significant differences in labelling were observed between OVA-OC-naïve and OVA-OC-experienced mice **(Figure S1C-S1H)**, suggesting that iEC-IEL crosstalk is not dramatically altered in food allergies. In contrast, we observed food allergy-dependent differences in mast cell and eosinophil interactions with iECs across the small intestine and colon. The number of both epithelium-associated and lamina propria (LP)-resident mast cells increased significantly in OVA-OC-experienced mice, and both mast cell populations exhibited substantial biotin labelling (**Figure 2B-2D)**. Unlike mast cells, the number of eosinophils increased significantly in the epithelium but not in the LP, and epithelium-associated eosinophils exhibited higher levels of biotin labelling than their LP counterparts. These data suggest that eosinophils exhibit pronounced epithelium-LP compartmentalization during food allergy (**Figure 2E–2G**).

Next, immune-epithelial contacts identified by uLIPSTIC were visualized using immunofluorescence imaging. Mast cells were rarely detected in OVA-OC-naïve mice, and in OVA-OC-experienced mice, they were found in close proximity to the epithelium and evenly distributed along the crypt-villus axis (**Figure 2H**). Eosinophils from OVA-OC-naïve mice were abundant in the lamina propria and did not exhibit crypt-villus bias. Following OVA-OC, eosinophils contacted the basolateral epithelial surface, and large epithelial-abutting eosinophil aggregates were found adjacent to the transit-amplifying and crypt regions (**Figure 2I**). Therefore, eosinophils reposition to interact directly with the stem cell niche after an allergic challenge.

To determine whether epithelial-interacting immune cells, and in particular eosinophils, have a distinct transcriptional identity, we performed single-cell RNA sequencing of FACS-purified, epithelial-associated immune cells from Villin-uLIPSTIC mice after OVA-OC. Cells were tagged with an anti-biotin oligo tag to quantify biotin labelling (Nakandakari-Higa et al., 2024; Stoeckius et al., 2018; Zheng et al., 2017). Unsupervised clustering identified five major immune cell lineages: mast cells, T cells, and pDCs displayed higher biotin labeling than eosinophils and ILC3s (**Figure 3A, Data Table 3**). Subclustering of mast cells revealed five subsets that exhibited identical levels of biotin labeling, indicating that transcriptionally distinct mast cell subpopulations interact equivalently with the epithelium in food allergies (**Figure 3B, Data Table 4**). In contrast, the four eosinophil subclusters differed significantly in biotin labelling (**Figures 3C, Data Table 5**). Subcluster-3, expressed genes (*Ccl9, Retnla*) associated with “basal” and “circulating” eosinophils (defined as per Gurtner et al., 2023), exhibited the lowest level of biotin labeling (**Figure S2A-2F**). Three “active” eosinophil subclusters were identified. Subcluster-0 and subcluster-1 displayed the highest levels of biotin labeling and were defined by wound repair (*Alox15, Emilin2*) and chemotaxis gene (*Dock2, Elmo1*) signatures, respectively (**Figure 3D-3E**). These biotin-hi, active eosinophil subclusters were enriched for secreted effectors (*IL-4 and Csf1*) and cell surface adhesion molecules (*Itgal and Alcam*) (**Figures 3F-3G, Data Table 5**). These data reveal that in food allergy, epithelial-interacting eosinophils adopt a transcriptionally distinct state enriched for wound repair, chemotaxis, and secretory programs, suggesting that eosinophils physically engaging the inflamed epithelium are poised to support repair.

**Figure 3.**
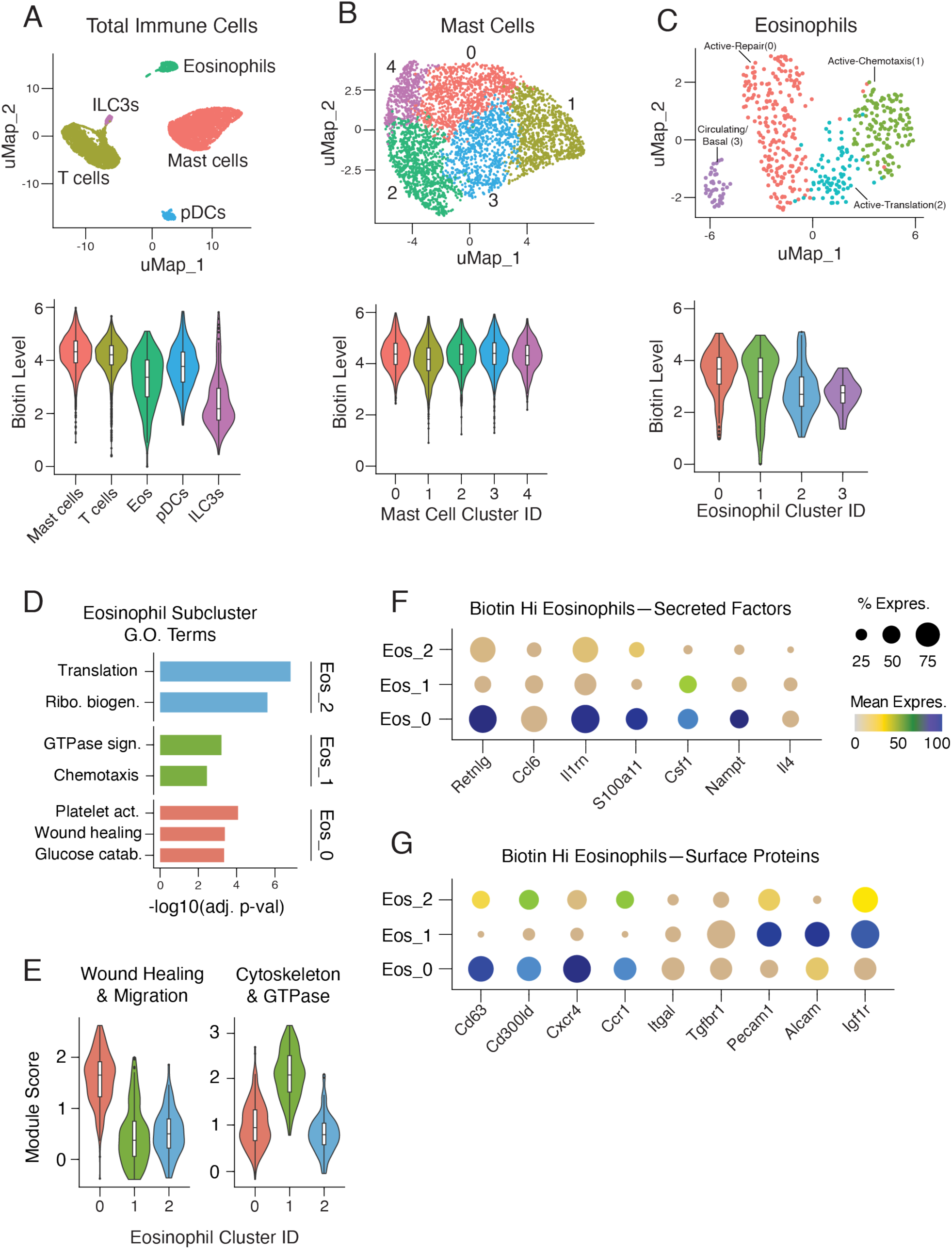
Epithelial-interacting immune cells have distinct transcriptional profiles. **A**. Single-cell RNA-seq analysis of immune cells isolated from the intestine of villin-uLIPSTIC mice exhibiting allergic diarrhea 3 h post sixth OVA-OC. Top, UMAP projection of gene expression-defined clusters identified using Seurat. Bottom: Corresponding biotin-oligo tag levels across clusters, identifying epithelial-interacting cell populations. **B**. Subclustering analysis of mast cells: Top, UMAP of gene expression-defined mast cell subclusters; Bottom, comparison of biotin-oligo levels between subclusters. **C**. Subclustering analysis of eosinophils: Top, UMAP of gene expression-defined eosinophil subclusters; Bottom, comparison of biotin-oligo levels between subclusters. **D-E**. Gene Ontology enrichment analysis of biotin-high eosinophil clusters (**D**) and corresponding functional module scores computed from genes identified in enriched pathways (**E**) using Seurat’s AddModuleScore function. **F-G**. Differentially expressed genes between the indicated eosinophil subclusters. The dot size represents the percentage of cells expressing each gene, and the dot color represents the expression level. For the violin plots throughout the figure, the boxes depict the interquartile range.

### Kinetics of iEC-innate immune interactions in food allergy

Allergic diarrhea and epithelial remodeling intensified with repeated OVA-OCs, suggesting that the underlying immune-epithelial interactions escalated over time. Therefore, we hypothesized that eosinophil-mast cell-iEC physical contact accumulates progressively with repeated allergen exposure. In the small intestine, the number of epithelial-associated eosinophils increased approximately ten-fold by three hours after each OVA-OC but returned to pre-challenge levels by 48 h. The colon displayed a different pattern: eosinophil numbers remained elevated after each challenge and accumulated progressively (**Figure 4A**). Despite robust early eosinophil recruitment, biotin labeling was minimal after the first OVA-OC and progressively increased after the fourth and sixth challenges, correlating with the severity of allergic diarrhea (τ=0.83, p=7.8e-9; **Figures 4B–4C** and **S3A–3B**). Minimal iEC labeling was observed in C57BL/6 mice subjected to the same protocol, confirming that robust eosinophil-epithelial interactions require oral food allergy susceptibility (**Figure 4B**). In contrast to eosinophils, the number of mast cells increased steadily with each oral challenge, and mast cell biotin labeling was stable across challenges, indicating constitutive rather than progressive iEC-mast cell interactions (**Figures 4D–4E** and **S3C-3D**). Immunofluorescence imaging revealed that eosinophil positioning shifted with successive challenges, with a relatively higher proportion of eosinophils localizing in the crypts at challenges 4 and 6. Large epithelial-abutting eosinophil aggregates formed transiently at these later challenges, consistent with the progressive biotin labelling observed by flow cytometry (**Figures 4F–4G**).

**Figure 4.**
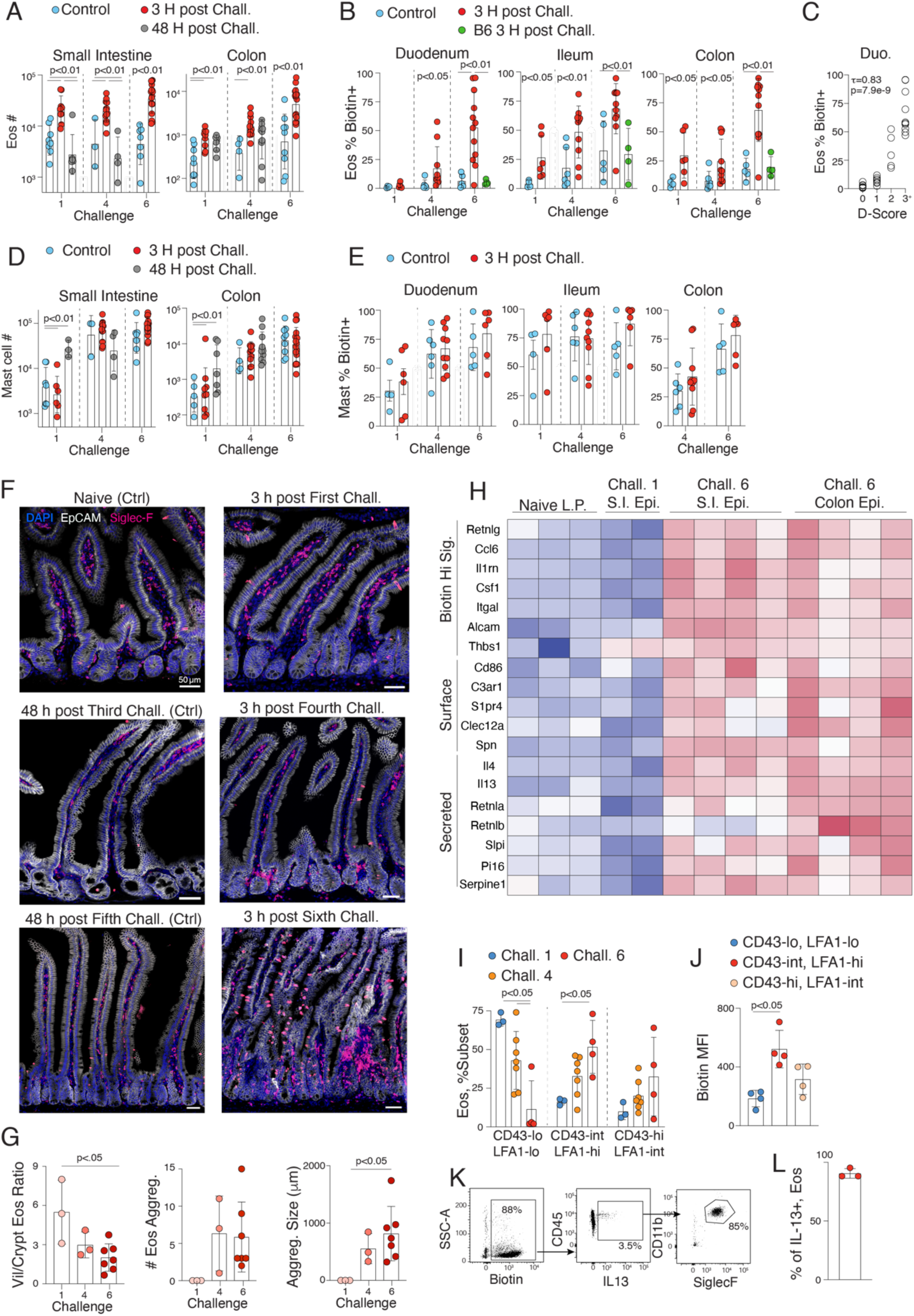
Eosinophil-epithelial interactions are kinetically coordinated with allergic challenges. **A-E**. Flow cytometry analysis of epithelium-associated eosinophils (CD45+, TCRγδ-, TCRβ-, CD11b+, SiglecF+) and mast cells (CD45+, TCRγδ-, TCRβ-, cKIT+, Fcer1+) at the indicated OVA-OC time points (1st, 4th, and 6th challenge). Controls for OVA-OC 4 and 6 received PBS gavage instead of the indicated challenge following three and five challenges, respectively. **A, D**. Quantification of the small intestine (left) and colon (right). **B, E**. Villin-uLIPSTIC biotin labeling in specific gut regions (duodenum, left; ileum, middle; colon, right) at the indicated challenge time points. **C**. Correlation analysis of diarrhea score with duodenal eosinophil-epithelial interactions using the Jonckheere–Terpstra test, τ=0.83, p=7.8e-9. **F-G**. Spatial and quantitative analysis of eosinophil positioning during allergic challenges. **F**. Representative images of the duodenum at indicated timepoints post OVA-OC (DAPI-dark blue, EpCAM-white, SiglecF-magenta). **G**. Quantification across 10 continuous crypts/villi: eosinophil position relative to crypt-villus axis (left), aggregate number (middle), and average aggregate size (right). **H**. Bulk RNA-seq analysis of FACS-purified eosinophils from the lamina propria of challenge-naïve mice (n=3), small intestine epithelium at first challenge (n=2), small intestine epithelium at sixth challenge (n=4), and colon epithelium (n=4). All genes displayed are significantly differentially expressed between conditions, Wald test with Benjamini–Hochberg FDR correction. **I-L**. Functional characterization of epithelial-interacting eosinophils. **I**. Flow cytometry quantification of CD43/LFA-1 expression subsets (CD43lo LFA-1lo, CD43int LFA-1hi, CD43hi LFA-1int) at indicated challenge timepoints. **J**. Comparison of Villin-uLIPSTIC biotin labeling at the 6th OVA-OC, stratified by the CD43/LFA-1 subset. **K-L**. Representative flow cytometry plots showing biotin labeling (left), IL-13 expression in biotin+ cells (middle), and quantification of eosinophils that are IL-13+ biotin+ (right). Data were log10-transformed, where applicable. One-way ANOVA with Tukey’s multiple comparisons or Welch’s T test was performed where applicable. Each dot in the bar graphs represents a biological replicate derived from a unique mouse, and the error bars represent one standard deviation from the mean. In heatmaps, each column represents a biological replicate derived from a unique mouse.

To characterize the transcriptional differences between eosinophils across OVA-OC time points, bulk RNA sequencing of FACS-purified eosinophil populations was performed. Relative to LP eosinophils from naïve mice, epithelial-associated eosinophils from sixth OVA-OC mice upregulated adhesion molecules (*Itgal, Spn*), chemotaxis receptors (*C3ar1, S1pr4*), and cytokines (*IL-4 and IL-13*) (**Figure 4H, S3E, Data Table 6**). Flow cytometry revealed that the majority of epithelial-associated eosinophils after the sixth OVA-OC were LFA-1^hi^ CD43^int^ (*Itgal* codes for the α-chain of LFA-1, *Spn* codes for CD43), and that these cells carried significantly more biotin than LFA-1^lo^ CD43^lo^ eosinophils (**Figures 4I–4J, S3F–3G**). Eosinophils accounted for the vast majority of IL-13-expressing biotin-positive cells after the sixth OVA-OC (**Figures 4K–4L** and **S3H**). This positions eosinophils as a local source of IL-13 within the stem cell niche, suggesting a direct role in tuft cell expansion in food allergies.

### Immune cell interactions with iEC subsets in food allergy

Food allergy suppressed Paneth cell gene expression and reprogrammed the intestinal stem cell niche toward a fetal-repair state (*see* Figure 1B, 1E). The progressive localization of eosinophils to the crypt and the formation of large epithelial-abutting aggregates adjacent to this niche suggest that eosinophil-Paneth cell physical interactions may underlie these changes. To directly test this and to map immune interactions across iEC subsets more broadly, we generated iEC subset-specific uLIPSTIC mice using Cre drivers targeting tuft cells (*Pou2f3*^GFP-CreERT2^; McGinty et al., 2020), Paneth cells (*Defa6*^Cre^; Adolph et al., 2013), and enteroendocrine cells (EECs; *Neurod1*^CreERT2^; Aprea et al., 2014) (**Figures 5A–5B**). Under physiological conditions, Paneth cells interacted with immune cells at a significantly higher rate than EECs or tuft cells after normalization to cell abundance, whereas tuft cell interactions were exceedingly rare (**Figures 5C–5E**). IEL subset analysis revealed that EECs preferentially labeled γδ T cells and CD8αα αβ T cells, whereas Paneth cells preferentially labeled CD8αα αβ T cells in the duodenum and CD4 αβ T cells in the ileum (**Figures 5F–5H** and **S4A–E**). These data establish that immune interactions with the intestinal stem cell niche are immune subset-specific and regionally compartmentalized under physiological conditions.

**Figure 5.**
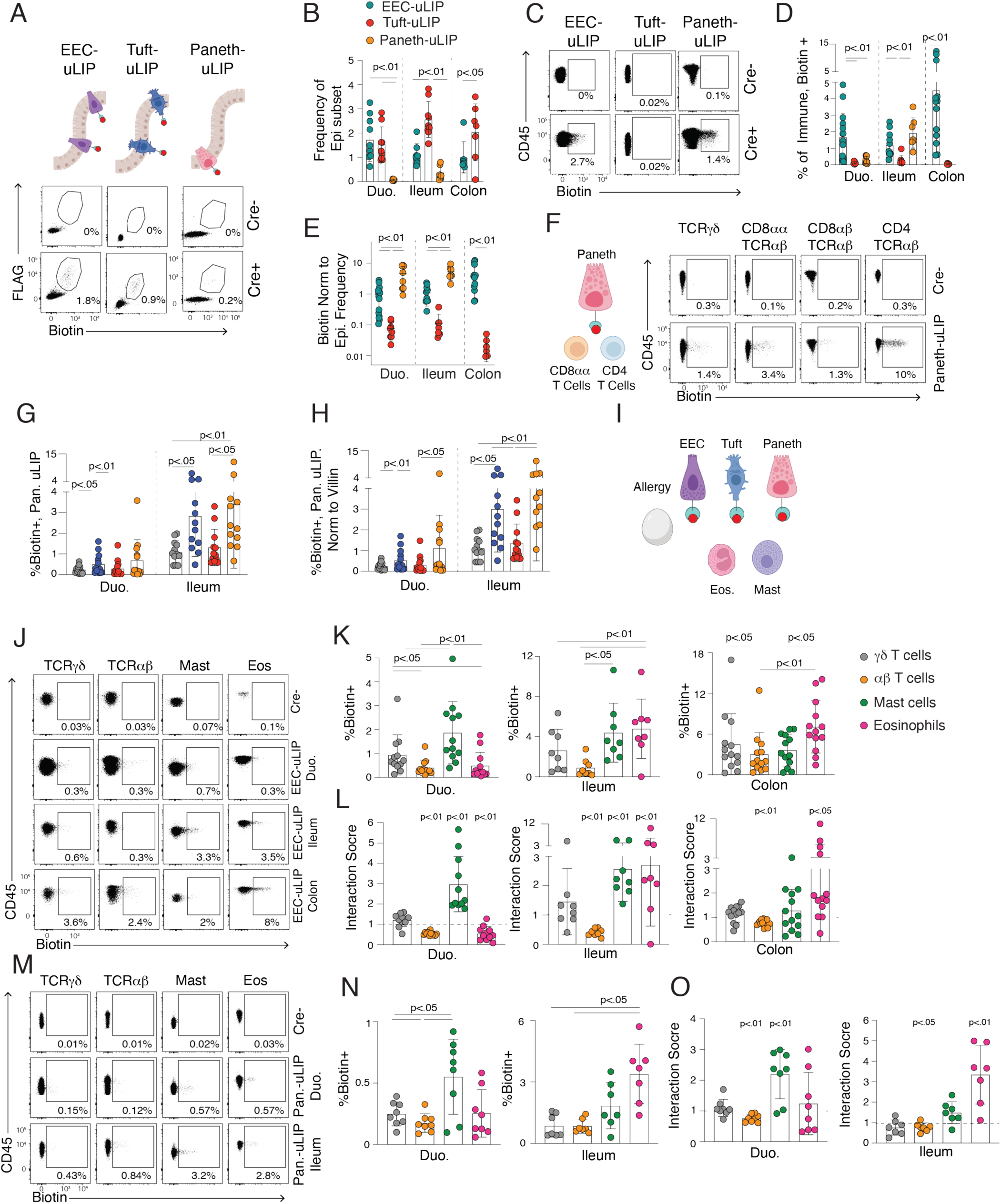
Epithelial subset-specific uLIPSTIC reveals compartmentalized immune-epithelial interactions. **A-H**. iEC subset uLIPSTIC analysis in the steady state. **A-B**. Quantification of FLAG+ (sortase-expressing) and Biotin+ epithelial cells (EpCAM+, CD45-) from Cre-negative control, Neurod1-CreERT2+ (enteroendocrine cell, EEC-uLIP), Pou2f3-CreERT2+ (tuft cell, Tuft-uLIP), and Defa6-Cre+ (Paneth cell, Panth-uLIP) uLIPSTIC mice. **A**. Representative FACS plots; **B**. Quantification of frequency for each donor iEC subset. **C-E**. Biotin+ immune cell (CD45+) labeling by each iEC subset uLIPSTIC strain. **C**. Representative FACS plots; **D**. Quantification of biotin+ immune cells; **E**. Normalization to FLAG+ epithelial cells. **F-H**. Characterization of T cell labeling across iEC subsets in the steady state. **F**. Representative FACS plots; **G**. Quantification of γδ T cells (CD45+, TCRγδ+, TCRβ-), CD8αα T cells (CD45+, TCRγδ-, TCRβ+, CD8α+, CD8β-, CD4-), CD8αβ T cells (CD45+, TCRγδ-, TCRβ+, CD8α+, CD8β+, CD4-), and CD4 T cells (CD45+, TCRγδ-, TCRβ+, CD8β-, CD4+); **H**. Normalization of the iEC subset uLIPSTIC to Villin-uLIPSTIC (see Supplemental Figure 4). I**-O**. iEC subset uLIPSTIC analysis following the sixth OVA-OC. **J, M**. Representative FACS plots; **K, N**. Quantification of biotin labeling of the indicated immune cell types (γδ T cells, αβ T cells, mast cells, and eosinophils). **L, O**. Interaction scores (calculated as: [% of biotin+ cells that are immune cell type A] / [% of biotin-cells that are immune cell type A]), indicating preferential engagement with specific iEC subsets. One-way ANOVA with Tukey’s multiple comparisons for all panels except L and O. For L and O, data were log10-transformed, and a one-sample T test against 0 was performed. Each dot in the bar graphs represents a biological replicate derived from a unique mouse, and the error bars represent one standard deviation from the mean.

Although tuft cells were both expanded and activated by the sixth OVA-OC, tuft cell-immune interactions remained minimal (**Figure S4F-G**). EECs exhibited substantial and regionally variable interactions following OVA-OC: mast cells were preferentially labeled in the duodenum, whereas eosinophils were preferentially labeled in the ileum and colon (**Figures 5J–5L** and **S4H–4J**). Paneth cells also displayed regional variable interactions: ileal eosinophils and duodenal mast cells were overrepresented among the interacting immune cells (interaction score > 1; **Figures 5M–5O** and **S4K–4M**). These data demonstrate that eosinophils physically engage with Paneth cells, providing evidence for direct eosinophil regulation of the stem cell niche.

### Eosinophils drive food allergy-associated epithelial remodeling

Eosinophil-Paneth cell contact at the stem cell niche, combined with eosinophils as the dominant local source of IL-13, suggests that eosinophils drive the epithelial remodeling observed in food allergy. To test this, we depleted eosinophils using two independent approaches in allergic mice (**Figures S5A-5F**). Anti-CCR3 targets eosinophils and basophils through opsonization, resulting in the efficient depletion of circulating and gut-resident eosinophils (**Figures 6A** and **S5A-5B;** Grimaldi et al., 1999). Anti-IL-5 antibody treatment blocks circulating IL-5, preventing eosinophil maturation and recruitment (Schumacher et al., 1988; Weng et al., 2011); depletion efficiency was variable across animals (**Figures 6A** and **S5C-5E**), enabling the correlation of depletion efficiency with functional readouts. Both cohorts were assessed for epithelial remodeling after four oral OVA challenges.

**Figure 6.**
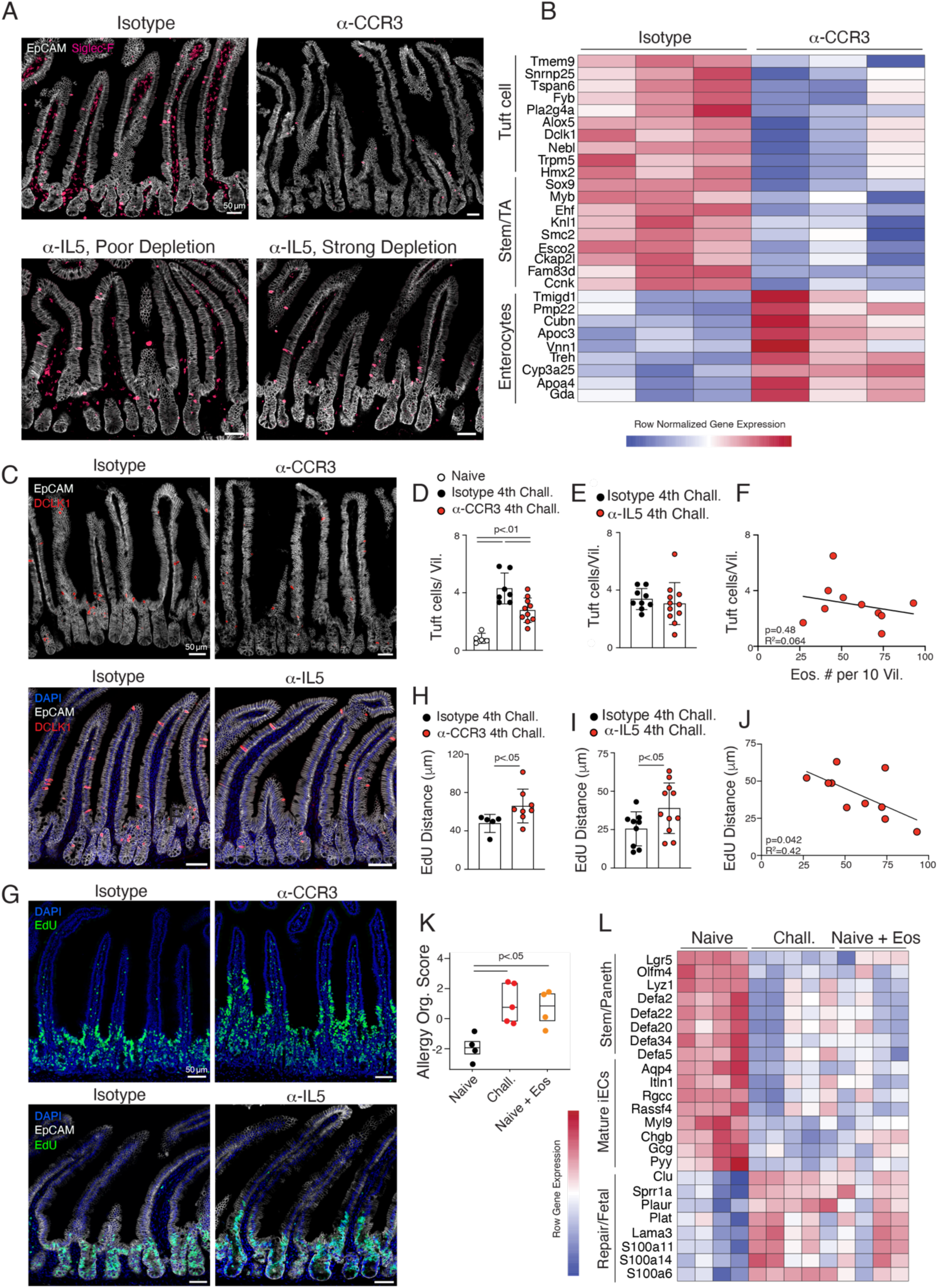
Eosinophils orchestrate allergen-induced epithelial remodeling. **A**. Representative images of the small intestine duodenum showing eosinophil depletion efficiency: isotype control (top left), anti-CCR3 treated (top right), anti-IL-5 treated with incomplete depletion (bottom left), and anti-IL5 treated with complete depletion (bottom right). **B**. Bulk RNA-seq analysis of FACS-sorted EpCAM+ small intestine epithelial cells from isotype control (n=3) and anti-CCR3 treated (n=3) mice 48 h post third OVA-OC. All genes displayed are significantly differentially expressed between conditions, Wald test with Benjamini–Hochberg FDR correction. **C-F**. Quantification of tuft cell abundance in eosinophil-depleted mice. **C**. Representative images (Dclk1-red, EpCAM-white, DAPI-blue). **D-E**. Tuft cell quantification per villus in anti-CCR3 (**D**) and anti-IL-5 (**E**) treated mice. **F**. Correlation between tuft cell number and remaining eosinophils in anti-IL-5 treated mice, linear regression analysis. **G-J**. Analysis of epithelial proliferation and migration in eosinophil-depleted mice. Mice were administered EdU 17 h prior to euthanasia and euthanized 48 h after the fourth OVA-OC. **H-I**. Quantification of EdU distance above the crypt-villus interface in anti-CCR3-(**H**) and anti-IL-5-(I) treated mice. **J**. Correlation between EdU migration distance and remaining eosinophils in anti-IL-5 treated mice, linear regression analysis. **K-L**. Organoid-based assessment of epithelial remodeling. Organoids were generated from challenge-naïve mice (n=4), OVA-OC challenged mice (n=5), and challenge-naïve mice co-cultured with epithelial eosinophils isolated 3 h after the sixth OVA-OC (n=4). An allergy organoid module score was derived from genes differentially expressed between control and allergy organoids and was compared across the three conditions. Heatmap displays genes significantly differentially expressed between conditions, Wald test with Benjamini–Hochberg FDR correction. One-way ANOVA with Tukey’s multiple comparisons for panel D; Welch’s T test for E, H, and I. Each dot in the bar graphs represents a biological replicate derived from a unique mouse, and error bars represent one standard deviation from the mean. For panels A, B, C, E, F, G, I, J, and K: mice were bred in-house at Rockefeller University. For panels D and H, mice were purchased from Jackson Laboratories.

Bulk RNA sequencing of epithelial cells isolated from control and anti-CCR3-treated allergic mice revealed a reversal of the OVA challenge-induced transcriptional program, including reduced tuft cell gene expression and a shift from a stem-like program to a mature enterocyte gene signature (**Figure 6B, Data Table 7**). Immunofluorescence imaging confirmed that anti-CCR3 treatment substantially reduced the tuft cell numbers per villus compared to isotype controls (**Figure 6C–6D**; for tuft cell counting in anti-CCR3 treatment, mice were sourced from Jackson Laboratories). Anti-IL-5 treatment did not reduce tuft cell numbers, possibly reflecting the incomplete eosinophil depletion achieved with this approach (**Figure 6E-6F**). Eosinophil depletion by either method restored epithelial proliferation and stem cell output as measured by EdU migration along the villus (**Figure 6G-6J**; for EDU measurement in anti-CCR3 treatment, mice sourced from Jackson Laboratories). In anti-IL-5 treated mice, residual eosinophil abundance was negatively correlated with EdU migration distance (r²=0.43, p=0.042; **Figures 6I–6J**), consistent with a dose-dependent relationship between eosinophil abundance and stem cell niche remodeling. These data establish that eosinophils are necessary for food allergy-associated epithelial remodeling, but leave open the question of whether this regulation is direct.

To test whether eosinophils are sufficient to drive epithelial remodeling independent of systemic allergic signals, we co-cultured intestinal organoids derived from challenged-naïve mice with eosinophils isolated from OVA-challenged mice. Eosinophil co-culture induced a transcriptional program resembling that of organoids derived from OVA-challenge-experienced mice, including the upregulation of fetal-repair genes (*Clu, Lama3*) and the downregulation of genes associated with mature enterocytes (*Aqp4*, *Itln1),* conventional stem cells (*Lgr5, Olfm4*), and Paneth cells (*Lyz1, Defa2)* (**Figure 6K-6L, Data Table 8**). These data establish that eosinophils are both necessary and sufficient to drive food allergy-associated intestinal epithelial remodeling, though whether this requires direct cell contact or is mediated by soluble factors remains to be determined.

### Eosinophil depletion exacerbates allergic diarrhea

The observation that eosinophils are required for food allergy-associated epithelial remodeling prompted us to investigate whether this remodeling is beneficial or detrimental to the host. To address this, we assessed the severity of allergic disease in eosinophil-depleted mice during repeated oral OVA challenges. In anti-CCR3-treated mice, the frequency and severity of allergic diarrhea were markedly increased compared to isotype controls across successive oral challenges, as measured by diarrhea incidence, diarrhea score (D-score) per challenge, and area under the D-score curve (**Figures 7A–7D**). Moreover, anti-CCR3-treated mice exhibited more severe acute body weight loss after challenge (**Figures 7E–7G**). Survival did not differ significantly between the isotype control and anti-CCR3 treated mice (**Figure 7H**). Anti-IL-5-treatment similarly increased the incidence and severity of diarrhea across oral challenges, with a significantly elevated cumulative D-score compared to the controls (**Figures 7I-7L**). Body weight loss was also more severe in anti-IL-5-treated mice than in controls (**Figures 7N–7P**). The efficiency of eosinophil depletion by anti-IL-5 treatment did not correlate with diarrhea scores or acute weight loss, suggesting that even partial eosinophil depletion is sufficient to exacerbate disease severity (**Figures 7M and 7Q**). Finally, anti-IL-5-treated mice showed a trend toward increased mortality following OVA-OC, consistent with anaphylactic shock (**Figure 7R**, p=0.053; for survival assessment in anti-IL-5-treatment mice sourced from Jackson Laboratory). Together, these data indicate that eosinophils play a protective role in food allergies by restraining the severity of allergic diarrhea and protecting against anaphylaxis.

**Figure 7.**
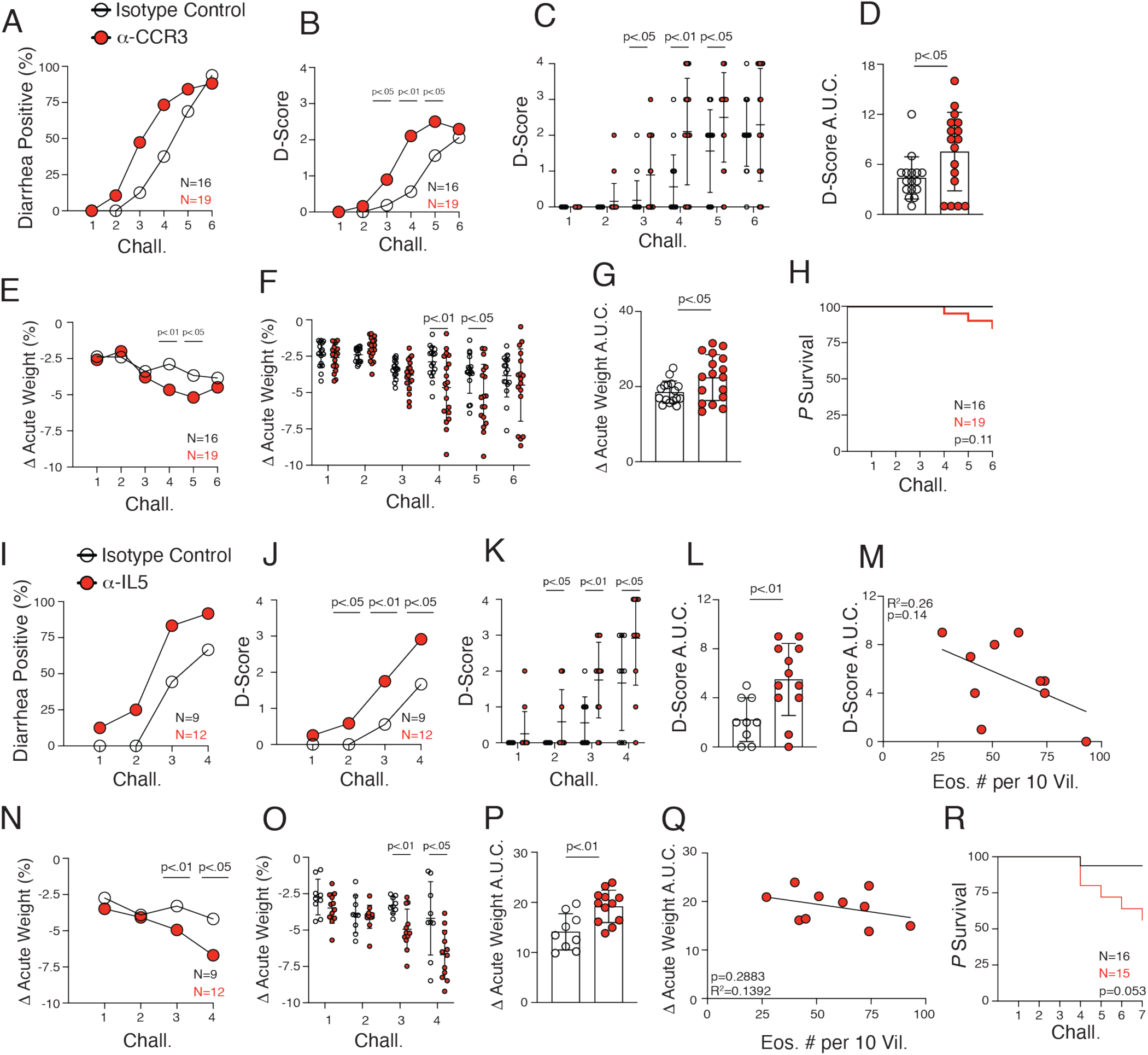
Eosinophil depletion exacerbates allergic phenotypes. **A-R**. Longitudinal assessment of allergic phenotypes in isotype control, anti-CCR3, and anti-IL-5 treated mice following OVA challenge. **A, I**. Diarrhea incidence. **B-D**, **J-L**. Diarrhea severity scored at each challenge; **D, L**. represent area under the curve. **E-G**, **M-P**. Body weight changes: acute weight loss (%) during the 2 h challenge window and area under the curve (**G and P**). **H, Q**. Correlation of diarrhea or acute weight loss with remaining eosinophils in IL-5 treated mice, linear regression. **R**. Kaplan-Meier survival curves. Student’s T test with Welch’s correction for B-G and J-P. Linear regression analysis for M and Q. Kaplan-Meier survival analysis for H and R. Each dot in the bar graphs represents a biological replicate derived from a unique mouse, and the error bars represent one standard deviation from the mean. For panels A-Q: mice were bred in-house at Rockefeller University. For panel R, mice were purchased from Jackson Laboratories.

## Discussion

This study demonstrates that eosinophils directly engage the intestinal epithelium to drive adaptive epithelial remodeling and restrain the severity of allergic diarrhea and anaphylaxis in mice. These findings suggest that eosinophils are critical regulators of mucosal homeostasis in food allergies rather than pathological effectors, consistent with a growing body of literature in murine models and patients (Ignacio et al., 2022; Rothenberg et al., 2024).

Eosinophils are tissue-resident granulocytes that populate the intestinal lamina propria and support the villus architecture (Ignacio et al., 2022; Mishra et al., 1999). Within the framework of a gut biological quality control system, eosinophils are established effector cells that mediate immunity to helminths and contribute to venom detoxification (Palm et al., 2012). Eosinophil infiltration and activation have been documented throughout the gastrointestinal tract in both food-allergic patients and mouse models (Hogan et al., 2001; Olbrich et al., 2020; Travers and Rothenberg, 2015). However, clinical trials targeting the eosinophil axis in gastrointestinal allergic diseases have failed to reproduce the therapeutic efficacy observed in patients with allergic asthma (Bel et al., 2014; Lugogo et al., 2016; Rothenberg et al., 2024). Our findings suggest that eosinophil-epithelial contact is protective rather than pathogenic, consistent with the failure of eosinophil-targeting therapies in allergic gastrointestinal diseases. Promoting eosinophil-epithelial engagement, rather than broadly suppressing eosinophils, may represent a more productive therapeutic direction, although the molecular determinants of this interaction remain unknown.

Similar to protective type 2 responses to helminths and protists (Gerbe et al., 2016; Howitt et al., 2016; von Moltke et al., 2016), food allergies drive tuft cell expansion. Eosinophils likely contribute to this expansion as the dominant local IL-13 source in the stem cell niche. In helminth infections, tuft cell-derived IL-25 amplifies type 2 immunity to accelerate parasite expulsion (Billipp et al., 2024; McGinty et al., 2020; von Moltke et al., 2016). Whether allergy-associated tuft cell expansion similarly amplifies protective responses or instead drives feed-forward pathology remains an open question. Overexpression of IL-25 exacerbates allergic diarrhea (Lee et al., 2016), raising the possibility that tuft cell expansion contributes to the severity of the disease. Tuft cell expansion is a component of a broader eosinophil-driven epithelial remodeling program that extends to the intestinal stem cell niche. Our results indicate that eosinophils are important modulators of this niche during food allergy, and their induction of a fetal-like repair transcriptional program suggests a role in limiting epithelial damage during recurrent allergen challenges.

The cell– and location-biases uncovered by subset-specific uLIPSTIC add cellular and regional resolution to our understanding of immune-epithelial crosstalk in physiology and food allergy. Paneth cell uLIPSTIC was of particular interest, as it revealed that the intestinal stem cell niche is a site of context-dependent immune regulation. In physiology, Paneth cells preferentially interact with CD4^+^ and CD8αα^+^ αβT cells, consistent with prior studies demonstrating MHCII-dependent T cell interactions with the stem cell niche (Biton et al., 2018). During allergy, this is supplanted by Paneth cell-eosinophil interactions that predominate in the ileum, providing a contact-dependent mechanism by which eosinophils regulate stem cell output and fate decisions (Quintero and Samuelson, 2025; Sato et al., 2011). Mast cell-Paneth cell interactions are abundant in the duodenum, reinforcing the role of this proximal segment as the primary site of allergen absorption and immediate hypersensitivity (Berin and Mayer, 2009; Noah et al., 2019; Reichel et al., 2026). Both physiology and allergy data suggest that immune regulation of the stem cell niche is spatially compartmentalized along the proximal-to-distal axis of the small intestine.

Beyond the stem cell niche, subset-specific uLIPSTIC uncovered immune-epithelial interactions with broad implications for intestinal physiology and allergic disease. In the steady state, EECs engaged IELs, extending prior evidence that EECs influence IEL gene expression through GLP-1 (Wong et al., 2022; Yusta et al., 2015), and raising the possibility that contact-dependent signaling also contributes to this crosstalk. During food allergy, EEC-eosinophil contacts predominated in the ileum and colon, suggesting that eosinophils may influence the allergic response through an unknown EEC-dependent mechanism (Gribble and Reimann, 2019; Worthington et al., 2018). Tuft cell-immune interactions were exceedingly rare across all conditions, consistent with the recently described capacity of tuft cells to evade CD8 T cell-mediated killing (Strine et al., 2024) and their function as a cellular reservoir for chronic mouse norovirus (Wilen et al., 2018), suggesting that tuft cells are broadly insulated from direct immune surveillance. Collectively, the iEC subset-specific uLIPSTIC tools developed in this study provide a generalizable framework for mapping immune cell interactions with defined epithelial subsets in any intestinal disease context.

Villin-uLIPSTIC enabled the identification of transcriptionally distinct eosinophil subsets with inherently different epithelial interaction propensities, revealing greater heterogeneity within the intestinal eosinophil compartment than previously appreciated. While Gurtner et al. classified gut eosinophils as “basal” or “active,” further subclustering was required to transcriptionally distinguish eosinophils with different epithelial interaction strengths within the active population. The epithelial-interacting LFA-1^hi^ CD43^int^ eosinophil subset is distinguished by the expression of immunomodulatory receptors (*Cd274, Tgfbr1, Tgfbr2*), adhesion molecules (*Itgal, Spn*), and cytokines (*IL-4, IL-13*), suggesting active regulation of local inflammatory cascades. Its gradual accumulation across successive allergen challenges is consistent with an adaptive role in shaping the mucosal response. Together, these findings establish a previously unappreciated protective function for eosinophils in food allergy, mediated through direct physical interactions with the intestinal epithelium, and define the cellular circuits by which eosinophils orchestrate mucosal adaptation to repeated allergen challenge.

## Limitations of the Study

Our study was conducted using the BALB/c OVA-alum food allergy model, which captures the key features of IgE-mediated food allergy but may not fully recapitulate the diversity of human food allergic responses across different allergens or genetic backgrounds. The eosinophil depletion strategies employed, anti-CCR3 and anti-IL-5, differ in their mechanisms of action. Key phenotypes (EdU migration and diarrhea severity) were observed across the approaches, providing strong evidence for an eosinophil-dependent mechanism. Differences in tuft cell expansion were observed between anti-CCR3 and anti-IL-5 treated mice. This may reflect the incomplete depletion of eosinophils by anti-IL-5 treatment. Additionally, while our organoid co-culture experiments demonstrated the sufficiency of eosinophils for epithelial remodeling, the precise molecular mediators of this interaction, including the role of IL-13, remain undefined. Finally, the extent to which the eosinophil-epithelial interactions and protective remodeling program identified here are conserved in human gastrointestinal food allergies awaits validation in patient samples.

## Author Contributions

P.W.D. and D.M. conceived the study. P.W.D., V.M.O., and Z.K. performed experiments. P.W.D and A.K. performed bioinformatic analysis. A.K. provided expertise in food allergy models. M.C.C.C and S.N.H. provided LIPSTIC expertise. G.V. provided uLIPSTIC reagents, mouse strains, and expertise. P.W.D. prepared the figures. P.W.D. and D.M. wrote the manuscript. All the authors have reviewed and edited the manuscript.

## Acknowledgements

We thank the members of the Mucida Laboratory for their helpful discussions. We are grateful to the Genomics, Flow Cytometry, Bio-Imaging Resource Center, Comparative Bioscience Cores, and the staff at The Rockefeller University for their support. We thank J. Aprea and F. Calegari for providing *Neurod1*^CreERT2^ mice, J.v. Moltke and I. Matsumoto for providing *Pou2F3*^CreERT2^ mice, K. Cadwell and R. Blumberg for providing *Defa6*^cre^ mice. We thank M. Hayashi and S. Liberles for discussions regarding EEC specific Cre drivers. We are grateful to R. Medzhitov, E. Florsheim, and J. Cullen for discussions regarding food allergy. We thank C. Wilen, M. Strine, and A. Okten for discussions on tuft cells. This work was supported by R01DK093674, R01DK113375, P01AI179273, CZI Science and Food Allergy FARE/FASI Consortium (D.M.); The Howard Hughes Medical Institute and R01AI157137 (G.D.V. and D.M.).

## Notice of pre-existing conditions, requirements, and licenses for article submission

The article submitted together with this notice is subject to the Immediate Access to Research policy of the Howard Hughes Medical Institute (“HHMI”). In accordance with this policy: (i) a preprint of this article either has been, or will be, deposited on a preprint server under a Creative Commons Attribution 4.0 International (CC BY 4.0) license and (ii) an additional author-published revised version of this article incorporating peer review feedback and/or new results or analysis either has been, or prior to journal publication will be, deposited on a preprint server under a CC BY 4.0 license. In addition, a non-exclusive CC BY 4.0 license to this article has been granted to the public and HHMI has a sublicensable, non-exclusive license to this article. This article is submitted for review and acceptance subject to these pre-existing conditions, requirements and licenses. If you have any concerns with any of these pre-existing conditions, requirements or licenses, please contact the corresponding author immediately

## Declaration of generative AI and AI-assisted technologies in the manuscript preparation process

During the preparation of this work, the authors used Claude for grammatical corrections and text editing. After using this tool/service, the authors reviewed and edited the content as needed and take full responsibility for the content of the published article.

## Methods

### Mice

C57BL/6J (000664), BALB/cj (000651), and B6.Cg-Tg(Vil1-cre/ERT2)23Syr/J mice (020282) were purchased from Jackson Laboratories and maintained in our facilities. ROSA26^uLIPSTIC^ mice were generated by G. Victora (Nakandakari-Higa et al., 2024). C57BL/6J-Tg(Neurod1-cre/ERT2)M1Fcal/J (025867) mice (Aprea et al., 2014) were kindly provided by J. Aprea and F. Calegari. B6(129S4)-*Pou2f3^tm1.1(cre/ERT2)Imt^*/J (037511) mice (McGinty et al., 2020) were kindly provided by J.v. Moltke and I. Matsumoto. Defa6-cre mice (Adolph et al., 2013) were kindly provided by R.S. Blumberg and K. Cadwell. All strains were backcrossed to the BALB/cj background for at least five generations and confirmed to be at least 90% BALB/cj homozygous through Transnetyx genetic monitoring before experimentation. All strains were backcrossed for at least eight generations before submission. Genotyping was performed in-house or by Transnetyx according to the protocols established for the respective strains by The Jackson Laboratories or personal communication with the donating investigators. Mice were maintained at the Rockefeller University animal facilities under specific pathogen-free conditions. Mice were fed a standard chow diet and used at 7–12 weeks of age for most of the experiments. Animal care and experimentation were consistent with NIH guidelines and were approved by the Institutional Animal Care and Use Committee at the Rockefeller University.

### Allergic Sensitization

Starting at 5 weeks of age, mice were sensitized by intraperitoneal (I.P.) injection of 50 μg of albumin from chicken egg white (OVA) (Sigma-Aldrich A5503-50G) in 0.2 aluminum hydroxide gel (Invitrogen, 21645-51-2), once a week for three consecutive weeks.

### Oral Challenge

At least 1 week after the final sensitization, the mice were gavaged with 75 mg of OVA in PBS on alternating days. For all experiments, mice were euthanized either 3 or 48 h after the indicated challenge, as detailed in the figure/figure legend.

### Diarrhea Tracking, Scoring, and Acute Weight Change

Mice were weighed before and 2 h after oral gavage with OVA to determine acute weight changes. During the 2 h window, mice were assessed every 15 min for diarrhea. Diarrhea score was assigned as follows: 0-no liquid stool observed at any assessment, 1-liquid stool observed at one assessment timepoint, 2-liquid stool observed at 2 assessment timepoints with minimal staining of the fur or tail, 3-liquid stool observed at 2 or more assessment timepoints with some staining of the fur and tail, 4-liquid stool observed at 3 or more assessment timepoints with extensive staining of the fur and tail.

### FACs purification of Intestinal Epithelial cells

The proximal half of the small intestine was removed from the mice, opened longitudinally, washed with PBS, and cut into 0.5 cm pieces. Gut pieces were shaken horizontally in a PBS solution with 5 mM dithiothreitol (DTT) (Fisher Scientific, R0862) for 10 min. The supernatant was discarded, and the gut pieces were suspended in RPMI-1640, 2% fetal bovine serum, 25 mM HEPES, and 1 mM sodium pyruvate (Complete RPMI) with 5mM EDTA. The gut pieces were incubated in this solution for 10 min at room temperature with gentle rocking. The supernatant was retained and the tissue was discarded. Cells were washed with complete RPMI and stained for 20 min at 4 °C with (EpCAM_G8.8_bioleged118210, CD45_30F11_BD_564279, and Live-Dead NearIR dye_Thermofisher_L34975). EpCAM+, CD45-, and Live-Dead-cells were FACs sorted directly into extraction buffer, PicoPure RNA isolation kit (Thermofisher KIT0204).

### Intestinal Epithelial cell RNA Purification, cDNA library synthesis, and Sequencing

RNA was extracted from the intestinal epithelial cells using the PicoPure RNA isolation kit (Thermo Fisher, KIT0204) following the manufacturer’s instructions. Libraries were prepared using the NEBNEXT Single Cell/Low Input RNA Library Prep Kit for Illumina (NEB# E6420S/L) and samples were indexed using NEBNext Multiplex Oligos for ILLUMINA (NEB #E6440S/L). Sequencing was performed on an Illumina NextSeq2000, P2 flowcell 1×100 a read length.

### Bulk RNA-sequencing Alignment

For all bulk RNA-seq alignments, raw reads were aligned to the mouse reference genome (mm39/GRCm39) using Subread (v2.0.6) with default RNA-seq alignment parameters. Coordinate-sorted BAM files were generated, and read counts per gene were quantified using featureCounts with the UCSC mm39 refGene gene annotation (GTF). This approach was used for all bulk RNA-seq experiments.

### Analysis of Intestinal Bulk RNA-seq

Transcript abundance files were used in the DESeq2 and ggplot2 R packages for all downstream differential expression analyses and generation of heat maps and supplemental data tables. Haber et al. 2017. was used as a reference to define tuft cell, goblet cell, and Paneth cell-specific genes.

### Preparation of intestinal Swiss-rolls

5-10 cm of the proximal small intestine was dissected from the mice, flushed with PBS, opened longitudinally, and Swiss-rolled. Swiss rolls were fixed overnight at 4°C in 4% paraformaldehyde in PBS, washed with PBS, dehydrated in 30% sucrose overnight, and embedded in O.C.T. Compound (Fisherbrand, #23-730-571) and stored at –20°C. Blocks were sectioned into 20µm-thick slices onto glass slides (Fisherbrand, #12-550-15) using a Leica CM3050 cryostat and air-dried for 1 h at room temperature.

### EdU administration

For the measurement of epithelial proliferation, the mice were injected intraperitoneally with 500 µg of 5-ethynyl 2’-deoxyuridine (EdU) (Thermo Fisher, #A10044) 17 h before sacrifice.

### Immunofluorescence imaging

The sectioned tissues were encircled using a PAP Pen (Vector Laboratories, #H-4000) to form a hydrophobic barrier and then rehydrated in PBS for 2 min to wash away the O.C.T. Compound. Sections were incubated with 100µL of a PBS solution containing 0.3% Triton-X-100 (Millipore Sigma, #93443), 1% bovine serum albumin (Millipore Sigma, #A1933), 2% normal goat serum (Jackson ImmunoResearch, #005-000-121), and 100U/mL heparin porcine salt (Sigma-Aldrich, #H3393) for 1 hour at room temperature to block non-specific fluorescence. Tissues were stained overnight at 4°C with the following primary antibodies: 1:100 anti-Siglec-F (E50-2440)-Alexa Fluor® 647 (BD Pharmingen, #562680), 1:200 anti-CD326/EpCAM (G8.8)-Alexa Fluor® 488 or 594 (BioLegend, #118210 and #118222), 1:100 Rabbit anti-DCAMKL1 (Abcam, #ab31704), 1:100 Rat anti-mouse Mast Cell Protease 1 (Mcpt1; Biotechne MAB5146), 1:100 anti-CD90.1 (AF647_OX-7_Biolegend_202508), 1:100 anti-FLAG_Cell Signaling Technology_14793S). Unconjugated antibodies were detected using secondary antibodies, including anti-Rabbit or anti-Rat Alexa Fluor® 488 and 647, at a 1:200 dilution for 1 h at room temperature. Nuclei were stained with 1:1000 1mg/mL DAPI Solution (ThermoFisher, #62248), and goblet cells with Wheat Germ Agglutinin (WGA), Fluorescein (Vector Laboratories #FL-1021-5). EdU was visualized using the Click-iT Plus EdU Imaging Kit and Alexa Fluor™ 488 dye (Thermo Fisher, #C10637) according to the manufacturer’s instructions. After staining sections were mounted using Fluoromount-G™ Mounting Medium (Invitrogen™ #00-4958-02) and a No.1½ 24×50mm cover glass (Corning®, #2980-245).

Immunofluorescence images were acquired using a Zeiss LSM 980 inverted confocal microscope with a 20X/0.80 air-immersion objective (1 airy unit pinhole) at the Bio-Imaging Resource Center of Rockefeller University (RRID: SCR_017791). Images were captured using one track per fluorophore, using the excitation laser lines of 405, 488, 561, 594, and 639 nm, and detected using GaAsP photomultiplier (PMT) tubes. Tiles acquired from the regions were chosen blindly. In each experiment, the control and experimental groups were acquired using the same acquisition settings, including laser intensity, detector gain, and pixel size, for accurate quantification.

### Image quantification

Images were pre-processed in FIJI (ImageJ) by adjusting the brightness and contrast, merging channels, applying the Despeckle filter to reduce random noise, and exporting files in TIFF format. Eosinophil and tuft cell quantifications were performed using the FIJI ‘Point’ tool, with coordinates stored in the Region of Interest (ROI) Manager for each intestinal villus. Eosinophil relocation was assessed by measuring the distance of each eosinophil from the crypt/villus interface. For quantification of epithelial cell proliferation, villus length, EdU distance from the base of the crypt, and crypt depth were collected using the ‘Straight Line’ or ‘Freehand Line’ tools, stored in the ROI Manager, and measured using the ‘Measure’ tool. The distance traveled by EdU from the crypt-villus interface was calculated by subtracting the EdU-positive length measurement from the total crypt depth. Quantification was performed for 10 villi per mouse. Representative images were prepared using Adobe Photoshop and Adobe Illustrator, and statistical analyses and data visualization were performed using GraphPad Prism.

### Organoid Generation and morphology characterization

The proximal half of the small intestine was removed from challenge-naïve mice or mice that underwent at least four bouts of OVA oral challenge and exhibited allergic diarrhea. The tissue was opened longitudinally, and the villous epithelium was mechanically removed by scraping with a scalpel blade. The tissue was then cut into approximately 0.25 cm fragments and washed in PBS with manual agitation, after which the tissue fragments were transferred to fresh PBS. For crypt release, the tissue was resuspended in PBS supplemented with EDTA (5 mM final concentration) and incubated on ice for 5 min. The suspension was triturated 10 times with a 25 mL serological pipette, and the supernatant was discarded. The tissue was then transferred to fresh 25 mL PBS + 5 mM EDTA and incubated at room temperature for 15 min with rocking. Following incubation, the suspension was triturated four times, and the supernatant was discarded. Sequential crypt fractions were collected by resuspending the tissue in 10 mL PBS and pipetting vigorously four times per tube. Each fraction was passed through a 70 μm cell strainer and centrifuged at 200 × *g* for 3 min. The pellet was resuspended in 10 mL PBS and centrifuged at 200 × *g* for 3 min, and the supernatant was discarded.

The crypt pellet was gently resuspended in IntestiCult organoid growth medium (StemCell Technologies, 06005) using a serological pipette to minimize the mechanical disruption. An aliquot of 50 µL was evaluated using light microscopy to assess crypt morphology and density, with a target of 50–100 intact crypts per well. For plating, crypts were embedded in 200 µL of a 60% Matrigel (Corning 356255) and 40% Intesticult solution and covered with 300 µL Intesticult in a pre-warmed flat-bottom 48-well plate. The medium and Matrigel were replaced three days after seeding. Spheroid or budded morphologies were assessed 6 days after seeding the culture. Bright-field images were acquired using a Zeiss LSM 980 with a wide-field camera, 20X objective.

### Preparation of Organoids for Bulk RNA-seq

After 6 days of culture, the organoids were liberated from Matrigel, washed with PBS, resuspended in PBS, vigorously pipetted to break apart organoids, and then resuspended in extraction buffer (PicoPure RNA isolation kit). RNA was extracted from organoids using the PicoPure RNA isolation kit (Thermo Fisher Scientific, Waltham, MA, USA; KIT0204) following the manufacturer’s instructions. Libraries were prepared using the NEBNEXT Single Cell/Low Input RNA Library Prep Kit for Illumina (NEB# E6420S/L) and samples were indexed using NEBNext Multiplex Oligos for ILLUMINA (NEB #E6440S/L). Sequencing was performed on an Illumina NextSeq2000, P2 flowcell 1×100 bp read length.

### Analysis of Organoid Bulk RNA-seq

Alignment of the RNA-seq data was performed as described above. Transcript abundance files were used in the DESeq2 and ggplot2 R packages for all downstream differential expression analyses and generation of heat maps and supplemental data tables. Mustata et al. 2013, was used as a point of reference to define the conventional and fetal/repair organoid signatures. The allergy organoid module score was calculated as follows: for each sample, a score was computed using two gene sets derived from prior differential expression analysis: an upregulated set consisting of genes significantly upregulated in allergy organoids that overlapped with a published fetal intestinal spheroid gene signature (75 genes) and a downregulated set consisting of genes significantly downregulated in allergy organoids that overlapped with a Paneth cell gene signature (eight genes). Raw counts were normalized using DESeq2 size factors, log2-transformed (log2[counts + 1]), and Z-scored across all samples for each gene. The module score for each sample was calculated as: Score = mean(Z-scores of up-regulated genes) − mean(Z-scores of down-regulated genes).

### uLIPSTIC Experimental Design

Biotin-Ahx-LPETGS-NH2 (substrate) was purchased by custom order from LifeTein and reconstituted in PBS at a concentration of 20 mM. The reconstituted substrate was stored at –70°C and thawed for one-time use. Mice were administered the substrate by intraperitoneal injection nine times at 20 min intervals, and euthanized 20 min after the final injection. The first injection volume was 400 µL, injections 2-9 were 200 µL. When applicable, OVA-OC or PBS gavage was administered contemporaneously with the second IP injection of the substrate. For all uLIPSTIC experiments, a Cre-uLIPSTIC control mouse was included in each experiment. These control mice were administered substrates and OVA-OC to match the experimental mice.

For all uLIPSTIC experiments involving a CreERT2 mouse line, mice were administered 100 μL of 50mg/mL of tamoxifen (Sigma Aldrich T5648) in 90% corn oil (Sigma Aldrich C8267) 10% ethanol (Sigma Aldrich E7023), by gavage, 72 hours and 24 hours prior to experimentation.

### Isolation of immune cells from the epithelium and lamina propria for flow cytometry

Intestines were harvested, opened longitudinally, washed in PBS, cut into 4 cm pieces, and incubated in PBS with 1mM dithiothreitol for 10 minutes at room temperature with gentle rocking. The supernatant was discarded, and the tissue was resuspended in 10mL of complete RPMI and 30 mM EDTA and orbitally shaken for 10 min at 37 °C. Intraepithelial cells were then recovered from the supernatant. For Lamina Propria cell isolation, the tissue was further washed with PBS, resuspended in complete RPMI with 50 mg/mL collagenase VIII and 200 mg/mL DNAse, and incubated for 30 min at 37°C with orbital rotation. The liberated cells were then washed with PBS. Intraepithelial and lamina propria-isolated cells were resuspended in a 37.5% percol solution (Thermo Fisher, 45001747), underlaid with a 75% percol solution, and centrifuged at 2300 RPM for 30 min. Immune cells were isolated from the 37.5%-75% Percol interface.

### Immune cell flow cytometry staining

For cell surface markers, cells were stained for 20 min at 4°C. For intracellular cytokine staining, the cells were fixed in Cytofix/Cytoperm buffer (BD 554722) for 30 min at 4°C and stained at room temperature for 30 min in perm buffer. Flow cytometry data were acquired on a BD Symphony. The following antibodies were used and listed in the format of antigen_clone_company_catalog: FCER1_MAR-1_BD_751758, Siglec-F_E50-2440_BD_740956, CD45_30-F11_BD_564279, TCRgd_GL3_BD_562892, Biotin_Bio3-18E7_Miltenyi_130-113-288, CD11b_M1/70_BD_553354, CD117_2B8_BD_558163, TCRb_H57-597_BD_553172, CD43_REA840_Miltenyi_130-112-888, LFA-1_REA880_Biolegend_141011, IL-13_eBio13A_ThermoScientific_25-7133-80, CD8a_53-6.7_BD_612898, CD8b_Ly_3_Biolegend_126627, CD4_GK1.5_eBioscience_25-0041-81.

### FACS Sorting for Single cell RNA-seq

Two sorting strategies were applied. A. Total Immune cells, enriched for non-eosinophils, non-B-, and non-T-cells: Live-Dead-negative, CD45-positive, CD19-negative, TCRgd-low, TCRb-low, CD11b-Siglec-F-double_negative. B. Eosinophils: Live-Dead-negative, CD45-positive, CD19-negative, TCRγδ-negative, TCRb-negative, CD11b-psotive, Siglec-F-positive. The following Total-Seq-C antibodies (BioLegend) were used to index samples from different mice and gut regions (duodenum, ileum, and colon): C0301, C0302, C0303, C0304, C0305, C0306, C0307, C0308, C0309, C0310, C0311, C0312, C0313, C0314, and C0315. Total-Seq-C antibody C0436 was used to quantify biotin levels.

### Single Cell RNA-seq Library Preparation

Single-cell libraries were prepared using the Chromium GEM-X Single Cell 5’ v3 kit (10x Genomics). Sequencing was performed on an Illumina NovaSeq SP flowcell with 800 million reads.

### Single Cell RNA-seq Alignment and Analysis

Single-cell libraries were processed using Cell Ranger Multi. Raw reads were aligned with the mouse reference genome (mm39/GRCm39). Single-cell libraries were analyzed using Seurat. Gurtner et al. 2023, was used as a reference point to define eosinophil subset signatures.

### FACS Sorting Eosinophils for Bulk RNA-seq

Eosinophils were sorted from the small intestine lamina propria of challenge-naïve mice, small intestine epithelium three hours after the first oral challenge, small intestine epithelium three hours after the sixth oral challenge, and large intestine epithelium three hours after the sixth oral challenge. Eosinophils were sorted as follows: Live-Dead-negative, CD45-positive, CD19-negative, TCRgd-negative, TCRb-negative, cKIT-negative, CD11b-positive, and SiglecF-positive. The cells were sorted directly into the PicoPure RNA isolation extraction buffer.

### Eosinophil Bulk RNA-seq Library Prep and Sequencing

RNA extraction, library preparation, RNA-seq alignment, and analysis with Desq2 were performed as described above for intestinal epithelial cells and organoids.

### CCR3 Depletion, Rockefeller Mice

Starting at 5 weeks of age, mice were sensitized by intraperitoneal (I.P.) injection of 50 μg of OVA in 0.2 aluminum hydroxide gel, once a week for three consecutive weeks. One week after the third sensitization, the mice were treated on alternate days until the end of the experiment with either 100 μg IP injections of anti-CCR3 clone 6S2-19-4 (BioXcell) or isotype control LTF-2 (BioXcell).

### CCR3 Depletion, Jackson Lab Mice

Starting at 8 weeks of age, mice were sensitized by intraperitoneal (I.P.) injection of 50 μg of OVA in 0.2 aluminum hydroxide gel once a week for two consecutive weeks. One week after the second sensitization, the mice were treated on alternating days until the end of the experiment with either 100 μg IP injections of anti-CCR3 clone 6S2-19-4 or isotype control LTF-2.

### IL-5 Depletion, Rockefeller University Facilities mice

Starting at 5 weeks of age, mice were sensitized by intraperitoneal (I.P.) injection of 50 μg of OVA in 0.2 aluminum hydroxide gel, once a week for three consecutive weeks. One week after the third sensitization, mice were treated with either 150 µg IP injections of anti-IL5 clone TRFK5 (Bioxcell) or isotype control LTF-2. Mice received four total antibody injections every third day prior to the first oral challenge. During the challenge phase, the mice were treated with antibodies on alternating days, which did not overlap with the challenges.

### IL-5 Depletion, Jackson Lab Mice

Starting at 8 weeks of age, mice were sensitized by intraperitoneal (I.P.) injection of 50 μg of OVA in 0.2 aluminum hydroxide gel once a week for three consecutive weeks. One week after the third sensitization, mice were treated with either 150 µg IP injections of anti-IL5 clone TRFK5 or isotype control LTF-2. Mice received four total antibody injections every third day prior to the first oral challenge. During the challenge phase, the mice were treated with antibodies on alternating days, which did not overlap with the challenges.

### Eosinophil Organoid Co-Culture

Six days after seeding, naïve organoids were liberated from Matrigel and manually disrupted by pipetting. 25,000-50,000 eosinophils were FACS-purified from the intestinal epithelium of mice 3 h after the sixth OVA-OC. Eosinophils were plated with these disrupted organoids in a 60% Matrigel and 40% Intesticult solution, covered in 300 µL Intesticult in a pre-warmed flat-bottom 48-well plate. The top layer medium was replaced two days after seeding. After 4 days of co-culture, the organoids were liberated from the Matrigel. RNA isolation, bulk libraries, sequencing, and analysis were performed as described above for the organoids and intestinal epithelial cells.

## Biological Sex

For the experiments throughout the manuscript (**Figures 1-5, S1-S4, Data Tables 1-6**) both male and female mice were used. Eosinophil depletion experiments (**Figures 6-7, S5, Data Tables 7-8**) were performed with female mice.

## Statistical analysis

Significance levels are indicated in the figure. All data are presented as the mean ± SD. At least two independent experiments were performed in this study. All statistical tests used were two-tailed, except for the Enrichment Score statistical analysis. The experiments were not randomized, and no statistical methods were used to predetermine the sample size. Multivariate data were analyzed using one-way ANOVA and Tukey’s multiple comparisons. Comparisons between the two conditions were analyzed using the Student’s t-test with Welch’s correction. Correlations were measured using a simple linear regression. GraphPad PRISM version 11.0 and R 4.4.3 were used to generate graphs and statistics.

## Data resources

All software used is available online, either freely or from a commercial supplier. No new software was developed for this project.

## Reagent Availability

No new materials were produced in this study.

---

**Supplemental Figure 1.**
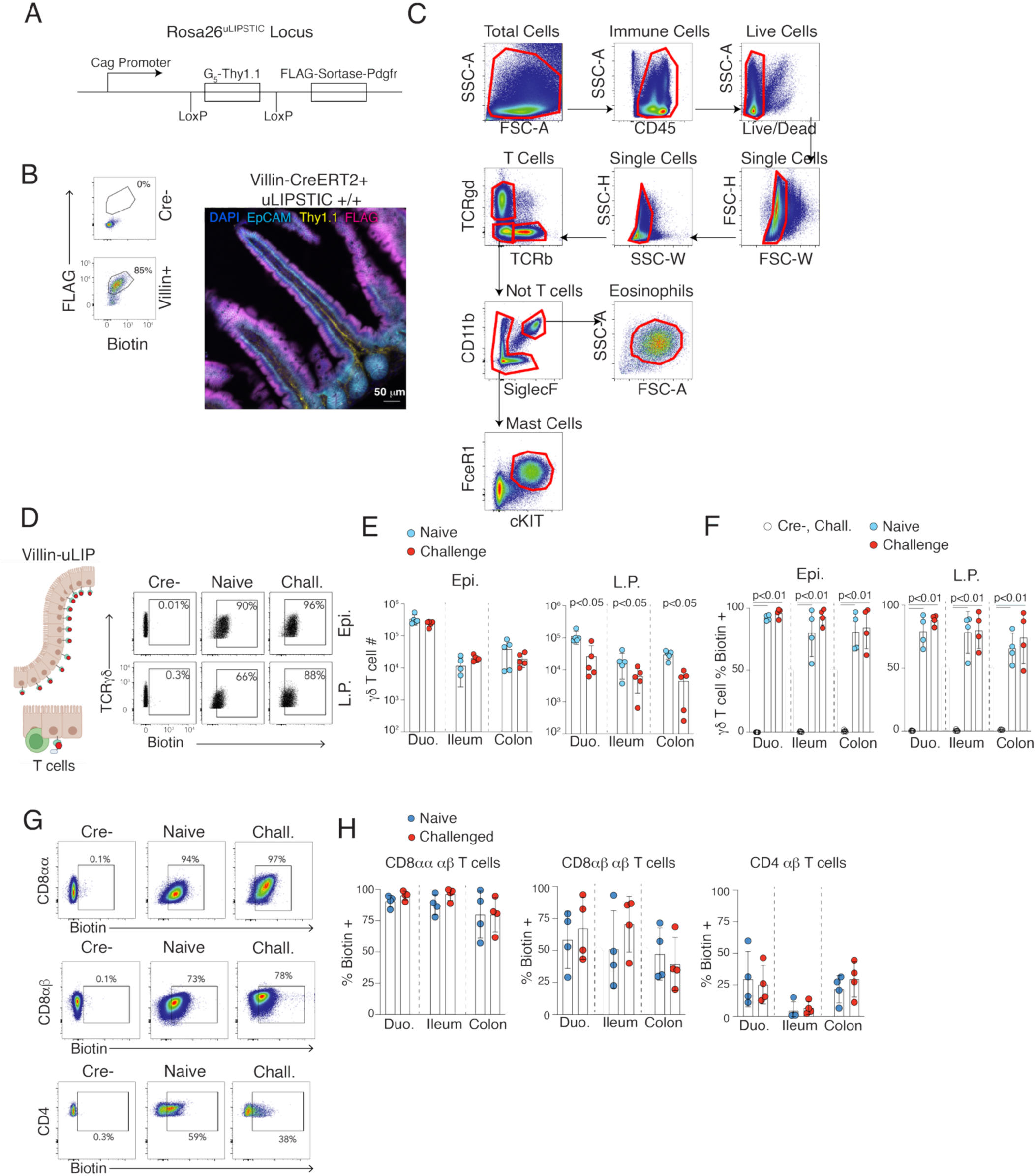
uLIPSTIC technology validation and gating strategies. **A**. Schematic representation of the Rosa26-uLIPSTIC allele, showing the genetic construct used for sortase-mediated labeling. **B**. Validation of Villin-uLIPSTIC functionality. Left: Flow cytometry staining of epithelial cells isolated from Cre– (negative control) and Villin-CreERT2+ uLIPSTIC +/+ mice for FLAG (tag bound to sortase) and Biotin (LPETG-Biotin substrate). Bottom: Representative confocal image of Villin-CreERT2+ uLIPSTIC +/+ epithelium (DAPI-dark blue, EpCAM-light blue, FLAG-magenta, Thy1.1-yellow). **C**. Flow cytometry gating schemes used throughout the study to identify γδ T cells (CD45+, TCRγδ+, TCRβ−), αβ T cells (CD45+, TCRγδ−, TCRβ+), mast cells (CD45+, TCRγδ−, TCRβ−, cKIT+, Fcer1+), and eosinophils (CD45+, TCRγδ−, TCRβ−, CD11b+, SiglecF+, SSC-Ahi). **D-H**. Flow cytometry analysis comparing biotin labeling between control and uLIPSTIC mice at steady state. **D-G**, Representative flow cytometry plots of biotin labeling in Cre-uLIPSTIC +/+ mice after the sixth OVA-OC (left), Villin-CreERT2+ uLIPSTIC +/+ challenge-naïve mice (middle), and Villin-CreERT2+ uLIPSTIC +/+ mice after the sixth OVA-OC (right), showing γδ T cells (**D**), CD8αα T cells (**E** top), CD8αβ T cells (**E** middle), and CD4 T cells (**E** bottom, shown in panel **G**). **H**, Summary data from n=4 mice per group (**E-H**). Welch’s T test was performed; no significant differences were observed between control and uLIPSTIC mice at baseline. Each dot in the bar graphs represents a biological replicate derived from a unique mouse, and the error bars represent the standard deviation.

**Supplemental Figure 2.**
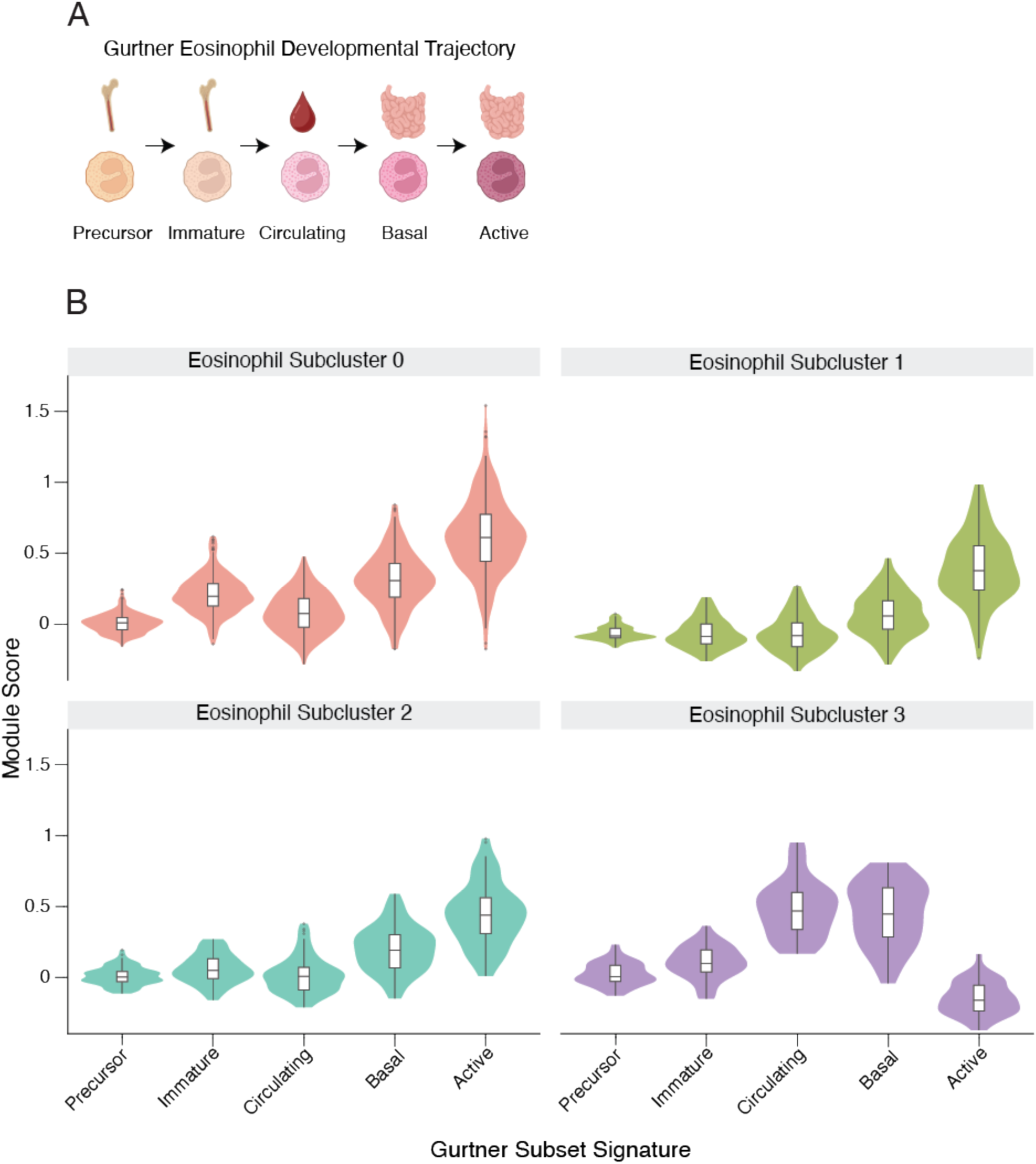
Eosinophil developmental state analysis. **A**. Schematic depicting the eosinophil developmental and differentiation trajectory proposed by Gurtner et al., illustrating the progression from immature to mature and active eosinophil states. **B**. Module score analysis of eosinophil developmental states. For each eosinophil cluster defined in Figure 3C, a gene signature corresponding to the EO_active population (derived from Gurtner et al.) was used to compute the module scores in the single-cell RNA-seq dataset. The gene lists were filtered to include only those genes that were detected in the dataset. Per-cell module scores were calculated using Seurat’s AddModuleScore function on log-normalized RNA expression data, which computes the average expression of the gene set subtracted from the expression of matched control gene sets to account for technical variation. Cells were grouped into Seurat-defined clusters, and module score distributions were visualized using violin plots. The module scores were summarized per cluster by calculating the mean, standard deviation, and cell count. Statistical differences in module scores across clusters were assessed using Kruskal-Wallis test, followed by pairwise Wilcoxon rank-sum tests with Benjamini-Hochberg correction for multiple comparisons.

**Supplemental Figure 3.**
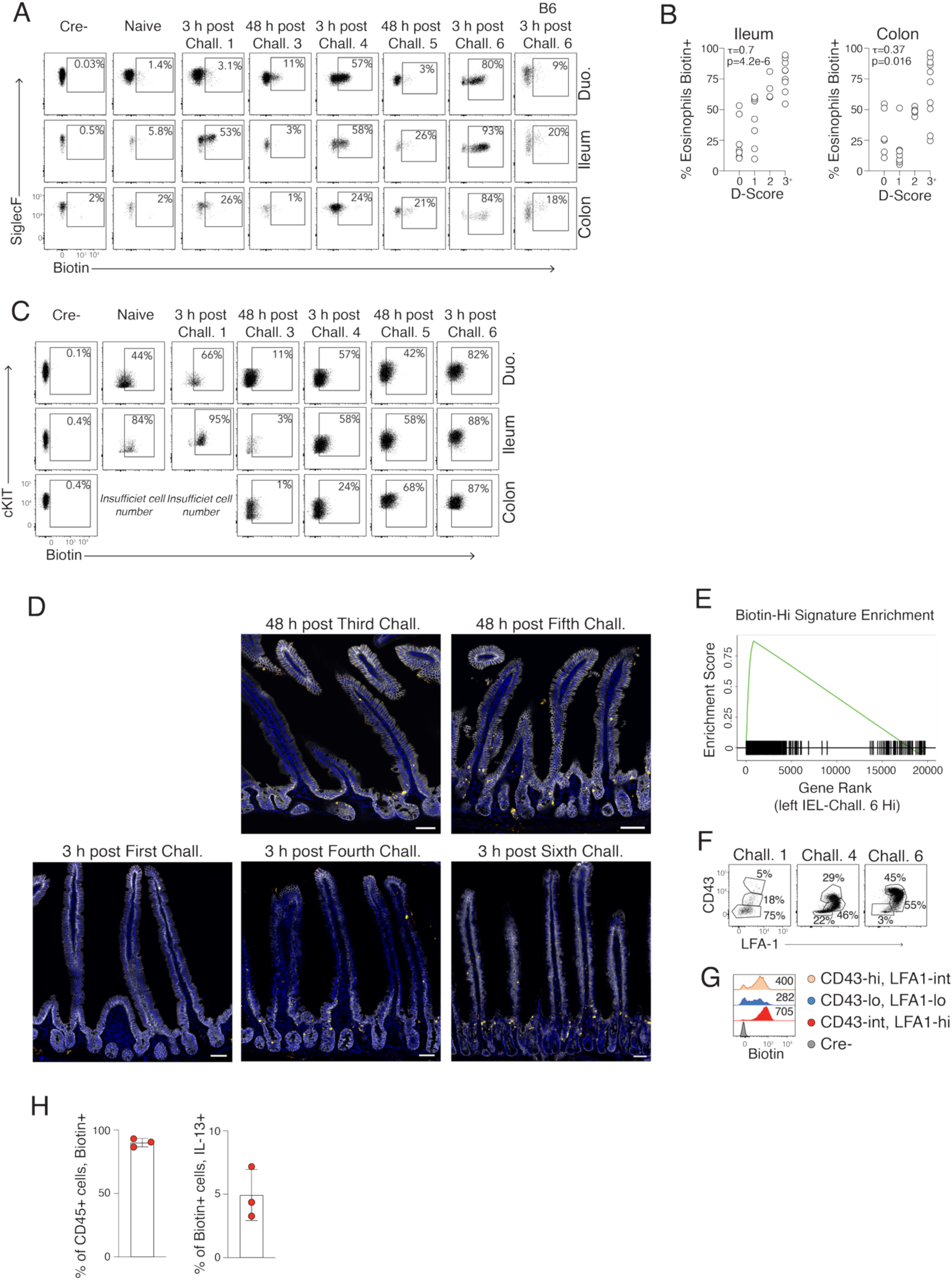
Kinetic and regional analyses of immune-epithelial interactions. **A**. Representative flow cytometry plots of Villin-uLIPSTIC biotin labeling of eosinophils (CD45+, TCRγδ-, TCRβ-, CD11b+, SiglecF+) at the indicated challenge time points. **B**. Correlation analysis of diarrhea score with Villin-uLIPSTIC biotin labeling of eosinophils in the ileum (left, τ=0.7, p=4.2e-6) and colon (right, τ=0.37, p=0.016). The Jonckheere-Terpstra test was performed. Each dot represents a biological replicate obtained from a unique mouse. **C**. Representative flow cytometry plots of Villin-uLIPSTIC biotin labeling of mast cells (CD45+, TCRγδ-, TCRβ-, c-KIT +, Fcer1+) at the indicated challenge time points. **D**. Confocal imaging of mast cells in the duodenum at the indicated challenge timepoints (DAPI-dark blue, EpCAM-white, Mcpt1-yellow). **E**. Gene set enrichment analysis comparing the Biotin-Hi eosinophil gene expression signature defined in the single-cell RNA-seq dataset to the bulk eosinophil RNA-seq dataset from Figure 4H. **F-G**. Functional characterization of eosinophil subsets. **F**. Representative flow cytometry plots depicting CD43 and LFA-1 staining of eosinophils at the indicated challenge time points. **G**. Villin-uLIPSTIC biotin labeling of indicated eosinophil subsets at sixth OVA-OC, stratified by CD43/LFA-1 expression. **H**. Quantification of Villin-uLIPSTIC biotin labeling. Left: Absolute number of biotin + immune cells. Right: Number of biotin+ IL-13+ cells. Each dot represents a biological replicate derived from a unique mouse, and the error bars represent the standard deviation.

**Supplemental Figure 4.**
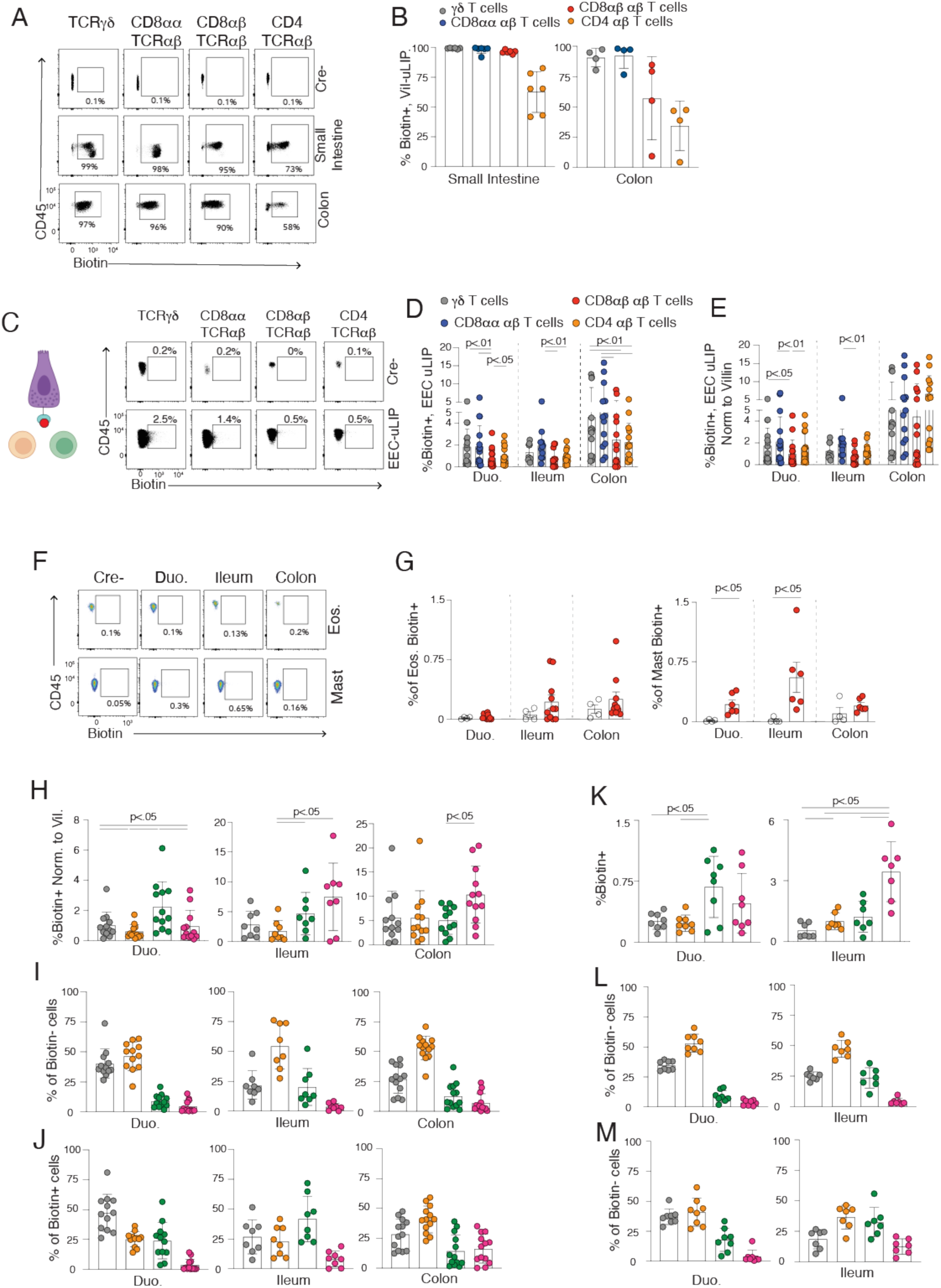
Epithelial subset-specific interactions in steady state and after challenge. **A-B**. Villin-uLIPSTIC analysis at the baseline. **A**. Representative FACS plots showing Biotin+ γδ T cells (CD45+, TCRγδ+, TCRβ-), CD8αα T cells (CD45+, TCRγδ-, TCRβ+, CD8α+, CD8β-, CD4-), CD8αβ T cells (CD45+, TCRγδ-, TCRβ+, CD8α+, CD8β+, CD4-), and CD4 T cells (CD45+, TCRγδ-, TCRβ+, CD8β-, CD4+). **B**. Quantification from n=4 mice. **C-E**. Neurod1-uLIPSTIC (enteroendocrine cell-specific) analysis at baseline. **C**. Representative flow cytometry plots. **D**. Quantification of Biotin+ immune cells. **E**. Normalization to Villin-uLIPSTIC data. **F-G**. Analysis of Pou2f3-uLIPSTIC (tuft cell-specific) after sixth OVA-OC. F. Representative FACS plot showing Biotin+ mast cells (CD45+, TCRγδ-, TCRβ-, Fcer1+, cKIT+) and eosinophils (CD45+, TCRγδ-, TCRβ-, CD11b+, SiglecF+). **G**. Quantification. **H-M**. Neurod1-uLIPSTIC and Defa6-uLIPSTIC (Paneth cell-specific) analysis after sixth OVA-OC. **H, K**. Biotin+ normalized to Villin-uLIPSTIC data for the indicated iEC subset uLIPSTIC strains. **I, J, L, M**. Percentage of total biotin-negative (**I, L**) or biotin-positive (**J, M**) cells that are the indicated immune cell type; these percentages were used to calculate the interaction scores shown in Figure 5L and 5O. T-test with Welch’s correction or one-way ANOVA with Tukey’s multiple comparisons correction was performed where applicable for panels D, E, G, and H-M. Each dot in the bar graphs represents a biological replicate derived from a unique mouse, and the error bars represent the standard deviation.

**Supplemental Figure 5.**
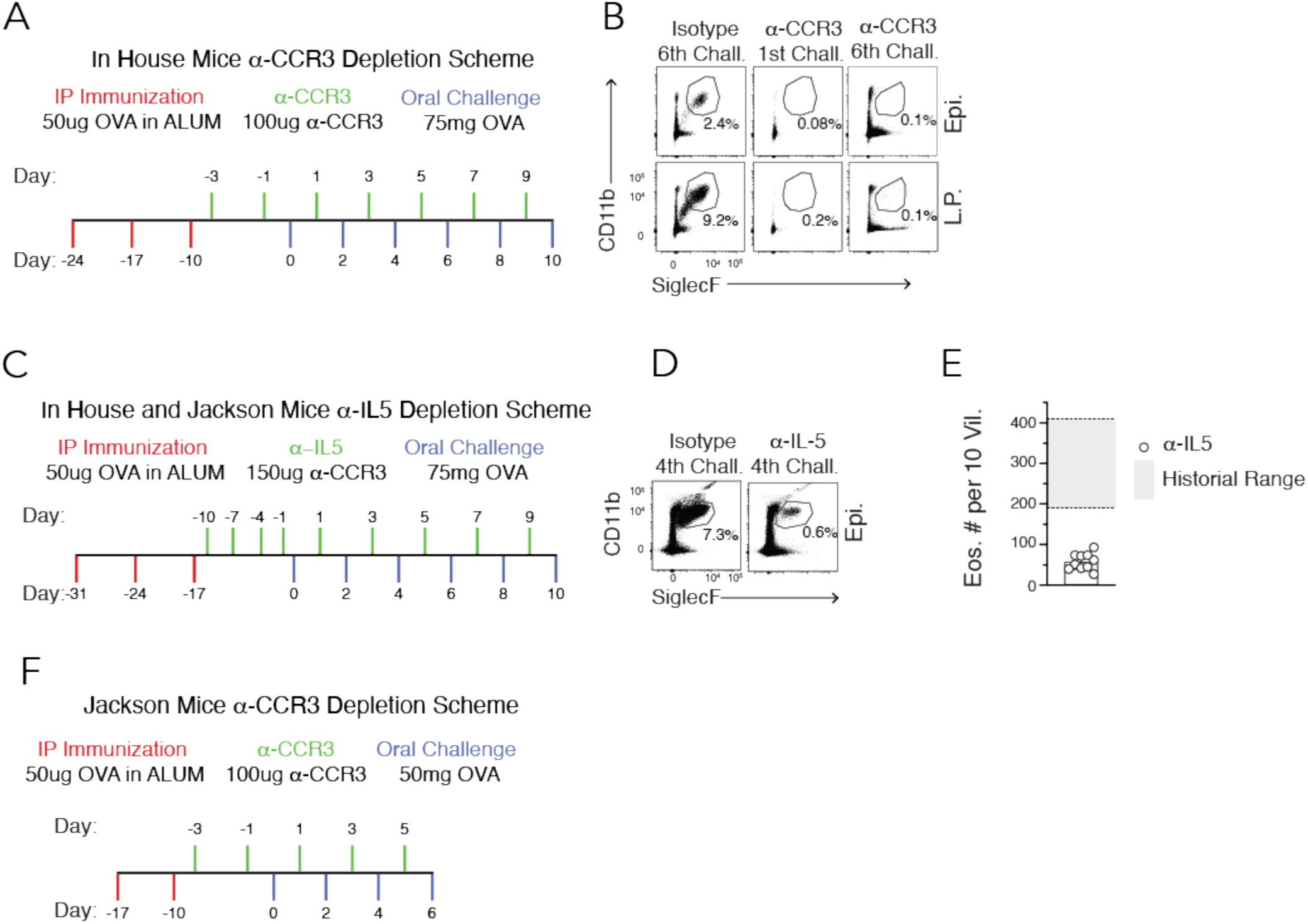
Eosinophil depletion validation and kinetics. **A, C, F**. Scheme representing of antibody treatment, sensitization, and oral challenge protocols for anti-CCR3 (**A, C**) and anti-IL-5 (**F**) eosinophil depletion approaches. Mice were sensitized by intraperitoneal injection of OVA-Alum weekly for three weeks and then treated with depleting antibodies prior to and throughout the challenge phase. **B, D**. Flow cytometry analysis of eosinophil depletion efficiency at the indicated time points using anti-CCR3 (**B**) and anti-IL-5 (**D**) treatments. Representative FACS plots showing biotin labeling of eosinophils in treated versus isotype control mice. **E**. Quantification of eosinophil numbers per 10 villi in anti-IL-5 treated mice at the indicated challenge timepoints, demonstrating dose-dependent depletion efficiency.

